# The composition of biophysical constraints generates complex, rugged regions of protein fitness landscapes

**DOI:** 10.1101/2025.04.12.648556

**Authors:** M. A. Spence, M. Sandhu, D. S. Matthews, J. Nichols, E. Stone, C. J. Jackson

## Abstract

The ruggedness of a protein’s fitness landscape constrains how it can evolve and how effectively it can be engineered. What determines whether a fitness landscape is rugged, and where in sequence space that ruggedness concentrates, remains poorly understood. Existing measures of ruggedness rely on complete combinatorial datasets and cannot be applied to landscapes in which only a fraction of possible sequences have been characterized. Here, we introduce a graph-based measure of ruggedness based on how quickly fitness differences diffuse across a landscape, which can be applied to sparse datasets. Applying this measure to 64 protein domains and several experimental systems with characterized folding and binding constraints, we show that ruggedness is not uniformly distributed but concentrates at regions where the dominant constraint on fitness changes. These results establish that the composition of individually simple constraints can generate localized ruggedness, providing a mechanistic basis for the complexity of protein fitness landscapes.

## Introduction

The topography of a fitness landscape dictates the dynamics of molecular evolution and constrains the efficacy of protein design^1–4^. For example, epistasis, where the phenotypic effect of a mutation is contingent on the genetic background^5–8^, generates “rugged” fitness landscapes. In such landscapes, small sequence changes produce non-linear, disproportionate changes in fitness^9^. In contrast, an underlying “smooth” fitness landscape implies that the effects of mutations are acquired gradually with mostly additive fitness consequences^9,10^. The ruggedness of a given protein landscape affects both natural evolutionary trajectories and the success of protein engineering campaigns^11–15^. Individual biophysical properties such as protein stability are often largely additive^16,17^, while composite phenotypes that depend on multiple constraints, such as enzymatic catalysis and ligand binding, typically show pervasive epistasis^18,19^. Enzymatic catalysis, for example, requires the concurrent maintenance of fold stability, substrate binding, transition state stabilization and product release. What determines whether a fitness landscape is rugged, and where in sequence space that ruggedness is concentrated, remains poorly understood.

Ruggedness has primarily been quantified through measuring epistasis across variants^16,20–27^, but this often requires that all combinations of mutations in a defined sequence space have been experimentally measured^24,28–30^. This limits its application to small combinatorial libraries and excludes the sparse datasets generated by experiments such as deep mutational scanning, directed evolution, *de novo* protein design and ancestral sequence reconstruction. Spectral landscape theory^31,32^ offers a solution to this, as it can decompose fitness variation into smooth and rugged components across a sequence graph and can be applied to incomplete, sparse datasets. In this framework, a rugged fitness landscape is one where fitness changes rapidly between sequences, while a smooth fitness landscape is characterized by gradual and correlated changes in fitness^33–37^. This can be formalized as a diffusion process, where fitness values spread between neighboring sequences in a graph, much like heat dissipating across a surface^38^. However, existing spectral graph approaches lack a single, comparable summary statistic that can be applied across diverse protein systems.

Here, we formalize this diffusion framework into a single measure of fitness landscape ruggedness and apply it to 64 protein domains assayed by deep mutational scanning under a common stability-based fitness function^39^. We find that fitness landscape ruggedness varies dramatically between domain folds and concentrates at mutations where the determinants of amino acid tolerance change, for example from hydrophobic core packing to surface polar interactions. We show that this arises because the nonlinear composition of selective constraints that separately generate smooth fitness landscapes necessarily produces ruggedness. This ruggedness concentrates where the limiting constraint changes between neighboring sequences, and we validate this mechanism across several experimental systems. These results establish that the composition of individually simple constraints can generate localized ruggedness, supporting a hypothesis that the complexity of a fitness landscape need not arise from the complexity of any single underlying constraint.

## Results

### A diffusion-based measure of ruggedness for sparse fitness landscapes

To study global structures in fitness landscapes, we first sought to define ruggedness as a numerical quantity that can be compared between landscapes, including sparse ones where most of the combinatorial sequence space has not been characterised. We represent each fitness landscape as a graph in which nodes are protein sequences and whose edges connect each sequence to its k nearest neighbours (kNN) in mutational distance, where mutational distance is the number of positions that two sequences differ (Hamming distance). When the data are dense enough, edges connect every pair of sequences that differ by a single point mutation. On such a graph, ruggedness is a spatial property that can then be quantified in terms of how much fitness changes along edges, and how far those changes propagate across the graph. Thus, a smooth fitness landscape would be one in which neighbouring sequences share similar fitness values and these similarities persist across many mutational steps; in contrast, a rugged fitness landscape would exhibit large fitness changes between neighbouring sequences.

To capture both how much fitness changes along edges and how far those changes propagate, we model fitness on the graph as a diffusion process: fitness values spread across the graph like heat dissipating across a surface. This is described by the heat diffusion kernel, *e^-tL^*, where *L* is the graph Laplacian and *t* is the diffusion scale, which controls how far fitness values are allowed to spread. (**SI Figure 1**). The heat diffusion kernel originates in the physics of heat conduction and has been adapted to graphs in spectral graph theory and machine learning, where it is a standard tool for describing how signals propagate across networks^38,40,41^. As *t* approaches 0, each sequence’s fitness is independent of its neighbors; as *t* approaches infinity all sequences converge on the mean fitness^38^ (**Figure 1a**). A larger value of *t* therefore corresponds to greater diffusion, stronger correlations in fitness between more distant sequences, and a smoother landscape.

**Figure 1.**
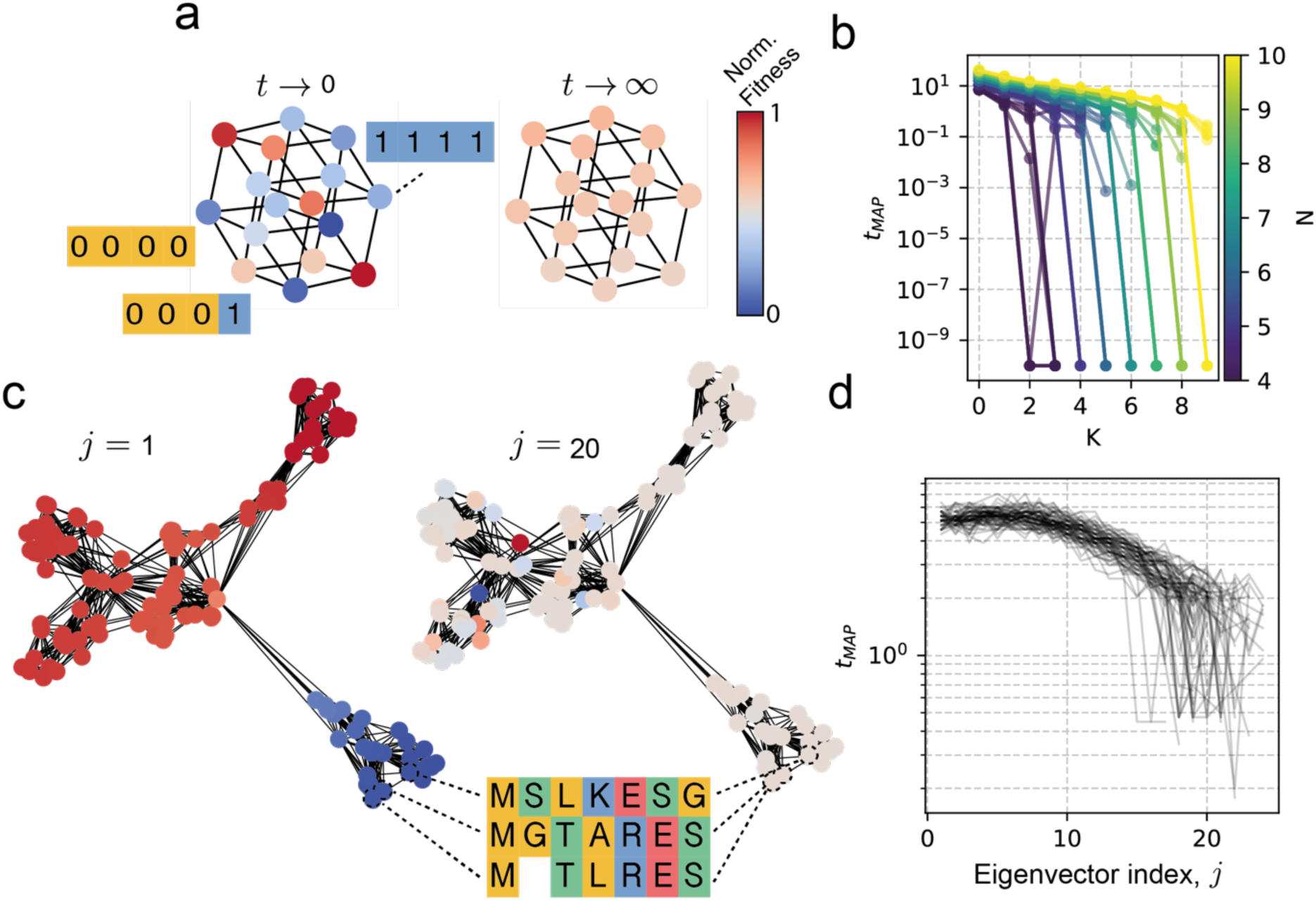
A diffusion-based measure of fitness landscape ruggedness. **a**. Heat diffusion on a fitness graph: at small diffusion scale *t*, fitness values remain local, whereas at large *t* they converge toward the graph mean. **b**. *t*_MAP_ decreases with the NK epistasis parameter K at fixed N. **c**. Sparse phylogenetic kNN landscapes whose fitness values are defined by low- and high-index Laplacian eigenvectors. **d**. *t*_MAP_ decreases with Laplacian eigenvector index on sparse simulated landscapes.

To estimate *t* from data, we need a model that links graphical structure to the observed fitness values. Here, we use a graph Gaussian process, which treats the fitness values across the graph as having a joint Gaussian distribution whose covariance structure is determined by *t*. We then find the value of *t* under which the observed fitness values are most likely. A single point estimate of *t* can be misleading, particularly for sparse landscapes where many values of t may fit the data nearly as well. We therefore use a Bayesian approach to compute a probability distribution over plausible values of *t* given the observed fitness data. We report the peak of this distribution, the maximum *a posteriori* value (*t*_MAP_), as our measure of ruggedness, where a larger diffusion scale implies stronger fitness correlations and thus a smoother fitness landscape.

Previous work has measured landscape ruggedness as the autocorrelation of a random walk over the graph^42–45^. *t*_MAP_, as presented here, and random walk autocorrelation are analytically related (**SI Figure 2**)^46^: both quantify the rate at which fitness is “forgotten” over sequence space. However, they differ in how they aggregate information across the graph. Random walk autocorrelation describes this decay separately for each mode of variation in the graph, whereas *t*_MAP_ describes it jointly over all modes at once, providing a single summary of global ruggedness.

This distinction is important for the kinds of graphs that arise from sparse fitness data, which are typically irregular and contain bottlenecks where most paths funnel through a narrow region of sequence space. On such graphs (such as “lollipop” and “dumbbell” topologies), random walk autocorrelation can oscillate or become negative, making the inferred correlation length unreliable. *t*_MAP_ is more robust to these features and converges to a well-defined lower bound under extreme ruggedness (**SI Figure 3**). On smooth graphs with high combinatorial completeness, the two measures broadly agree.

We validated this approach on two synthetic systems where landscape ruggedness can be controlled. The first of these is the NK-landscape model^47,48^, a standard test system in which each genotype is a sequence of N binary sites and the fitness of each site depends on its own state and the states of K other sites. K therefore tunes the amount of epistasis in the system: K = 0 gives a purely additive landscape, and increasing K introduces higher term interactions and more ruggedness. As expected, *t*_MAP_ correlates strongly and negatively with K at fixed values of N (Spearman’s ρ=-0.76; **Figure 1b**), confirming that *t*_MAP_ responds to controlled changes in epistasis. However, NK landscapes are dense and combinatorially complete, which does not match the structure of most real fitness data. To test *t*_MAP_ on sparse landscapes we simulated sequences along random phylogenetic trees^49^ and built kNN graphs from the resulting alignments. This produces a realistic sparse sequence space with irregular connectivity, rather than a complete combinatorial hypercube. We then defined fitness functions on these graphs using eigenvectors of the graph Laplacian. The Laplacian’s eigenvectors are the natural modes of variation on the graph, analogous to sine waves of different frequencies: low-index eigenvectors vary smoothly across the graph, and high-index eigenvectors oscillate more rapidly. Using these as fitness functions therefore generates landscapes whose ruggedness is set directly by the eigenvector index, without needing to specify any epistatic interactions^46,50^ (**Figure 1c**). *t*_MAP_ correlates strongly with the Laplacian eigenvector index (Spearman’s *ρ* = -0.94; **Figure 1d**), indicating that the measure works on sparse, irregularly connected systems in addition to dense combinatorial ones (such as NK).

Ruggedness, as we measure it here, is distinct from epistasis. Ruggedness is a global geometric property of the fitness landscape: it captures how far apart in sequence space two genotypes can be before their fitness values become uncorrelated. Epistasis, by contrast, is defined by the specific interaction terms in a decomposition of the genotype-phenotype map^24^. The two are related but not equivalent, and the distinction matters when comparing landscapes of different sizes (**SI Figure 4**). Even in the complete absence of epistasis (K=0), larger combinatorial sequence spaces appear more rugged by *t*_MAP_, because more eigenmodes of the graph Laplacian must be attenuated for fitness to decorrelate across the full space. We observe this directly in the NK landscapes, where *t*_MAP_ correlates with N at K=0 (**SI Figure 5**). The reason is that larger sequence spaces can host higher-order interaction terms that smaller ones cannot, and at a constant low degree of epistasis these terms remain unrealized, making the larger landscape appear smoother on a per-mutation basis. Indeed, normalizing the Laplacian eigenvector index by N abolishes this correlation (**SI Figure 6**), confirming that the apparent N dependence reflects the size of the spectrum rather than any change in underlying epistatic structure.

### When fitness is defined by stability, ruggedness varies by fold and concentrates where the limiting stabilizing interaction changes

Having established *t*_MAP_ as a measure of ruggedness, we next asked whether ruggedness varies between protein folds under identical selective pressure. To avoid confounding effects from differences in selective pressure over multiple studies, we used deep mutational scanning (DMS)^51^ data from a single large-scale experiment in which multiple protein domains were each assayed for thermodynamic stability under an identical unfolding selection^39^. We modeled the fitness landscapes of each domain as a Hamming graph where nodes are sequence variants and edges connect sequences differing by a single substitution. Across the 64 domains we tested^52^, *t*_MAP_ varied from 2.8 (most rugged) to 6.3 (smoothest) (**Figure 2a**). This indicates that ruggedness depends on the protein fold, even when the fitness function is constant. Individually, all observed landscapes were significantly smoother than the corresponding null models generated by random permutation of fitness values across nodes (individual p-values < 0.02, 100 permutations per system) (**Figure 2b, c**), confirming that the smoothness reflects genuine sequence-fitness structure rather than graph topology.

**Figure 2.**
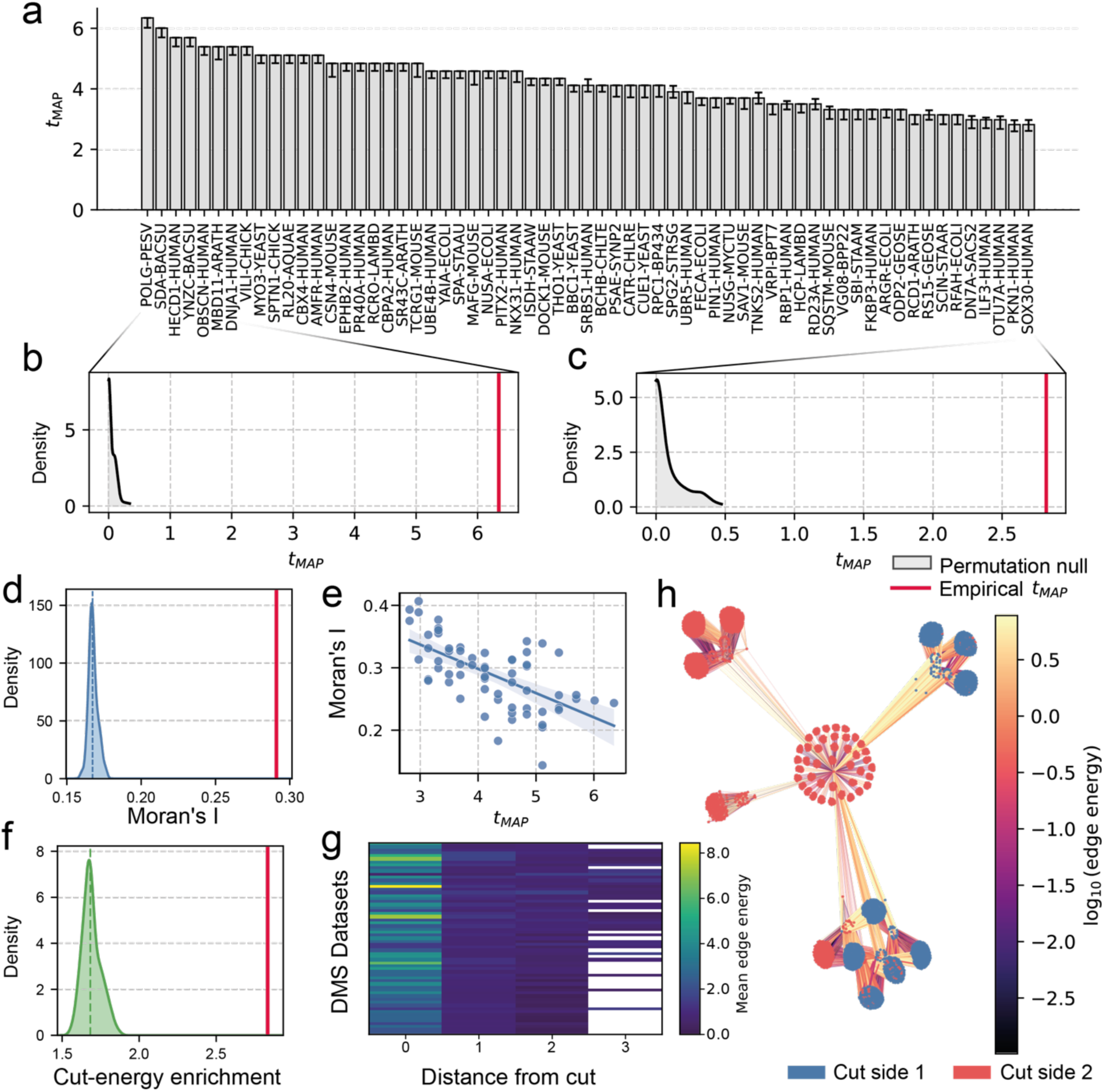
Fitness landscape ruggedness localises to high energy cutsets. **a**. *t*_MAP_ values across 64 protein domains assayed under a common stability-based unfolding selection. **b, c**. Node-permutation null distributions for *t*_MAP_ in the least and most rugged domains, with empirical values marked in red. **d**. Node-permutation null for Moran’s I of edge Dirichlet energies, with the empirical value marked in red. **e**. Spatial autocorrelation of edge energy decreases with *t*_MAP_ across domains. **f**. Node-permutation null for spectral cut-energy enrichment, with the empirical median enrichment marked in red. **g**. Mean edge energy decays with geodesic distance from the spectral cutset across DMS datasets. **h**. Example spectral bipartition of POLG-PESV, with nodes coloured by cut side and edges coloured by Dirichlet energy.

We next asked whether the ruggedness observed in these landscapes is diffuse across sequence space or concentrated in specific regions of it. To quantify local fitness variation (i.e., ruggedness) we computed the Dirichlet energy of each edge, defined as the squared fitness difference between the two connected sequences. An edge with a large fitness change between its endpoints, characteristic of a locally rugged region of the landscape, has a high Dirichlet energy. We treated high-energy edges as a graph-localized signal and tested spatial organization directly by autocorrelation. Indeed, high energy edges share a spatial autocorrelation (Moran’s I = 0.291, Geary’s C = 0.722) that is significantly greater than expected under a node permutation null (p-value < 0.01; 100 permutations; **Figure 2d, SI Figure 7**). As expected, the strength of the spatial autocorrelation decreases with *t*_MAP_ (Spearman’s Rho = 0.62; **Figure 2e, SI Figure 7**), indicating that ruggedness is locally concentrated along spatially coherent mutations in rugged landscapes. We tested this further by by applying spectral bipartitioning to each Hamming graph, weighting the edges by their inverse Dirichlet energy so that the resulting partition preferentially passes through high-energy edges (**Methods**) while remaining a spectrally balanced cut (**Figure 2h**). In 62 of 64 domains, edges along the cut carried significantly more fitness variation than edges within either subgraph (median enrichment = 2.83-fold; **Figure 2f**), and aggregate edge energy decayed with graph geodesic distance from the cut (**Figure 2g)**. Because the spectral partition is weighted by per-edge Dirichlet energies (**Methods**), the location of the cut depends on the fitness signal in addition to the graph topology. To verify this, we re-ran the bipartitioning on each Hamming graph after randomly permuting fitness values across nodes (preserving topology); the cutset enrichment collapsed in all 64 domains (median enrichment 1.68-fold across systems, individual p-values < 0.01, 100 permutations per system; **SI Figure 8**), confirming that both the spatial localisation of ruggedness reflects fitness structure rather than the topology of the Hamming graph and that under the node-wise permutation null distribution, Dirichlet energy is not enriched along an energy enriched spectral cutset. These results together show that ruggedness concentrates at a spatial boundary and is not broadly diffuse across sequence space.

Given that ruggedness localizes to specific regions of sequence space, we next asked what determines where these boundaries occur. Thermodynamic stability depends upon several distinct biophysical forces, including hydrophobic packing, hydrogen bonding, backbone geometry, and solvation, with different folds weighting these contributions differently^39,53–55^. The mutations a protein can tolerate therefore depend on which force is currently most limiting; a sequence whose stability is closest to failing through poor core packing will tolerate different substitutions than one limited by surface solvent interactions. We hypothesized that ruggedness concentrates at the boundaries between regions of sequence space dominated by different stabilizing constraints, where the rules governing tolerated substitutions can change abruptly. Indeed, the number of independent traits under selection has long been recognized as a determinant of landscape complexity^56–58^.

To test this, we needed a way to estimate the biophysical constraints that dominate fitness effects at each position in a sequence from DMS data. For each domain, we built a mutation effect matrix **M**, with one column per sequence position and one row per amino acid, where each element records the fitness effect of the specific substitution at that position (**Figure 3a**), and decomposed **M** using singular value decomposition (SVD)^59^. SVD identifies the dominant patterns of covariation between positions and amino acids and ranks them by how much of the variance in **M** each pattern explains (**Figure 3b**). Each SVD axis groups together positions that share the same or similar patterns of tolerated substitutions. If a single pattern in **M** governed fitness effects across the whole fold, one axis would dominate; additional axes may have higher loadings when subsets of positions follow systematically different patterns of fitness effects. We thus treated each SVD axis as a proxy for a distinct biophysical constraint acting on the fold, an approach used previously to extract functional constraints from DMS and coevolution data^60–62^. Importantly, SVD axes are not absolute or interpretable biophysical constraints, but are scale-dependent groupings of positions that share a common pattern of tolerated substitution that we treat as a data-driven indicator of a biological constraint. Two observations support this interpretation. First, positions belonging to the same axis are spatially clustered in the domain 3-dimensional structure beyond expected under a null permutation (50/64 domains, on each domain p-values < 0.05, 300 permutations per system; **SI Figure 9**). Second, the preferred substitutions differ systematically between axes (**SI Figure 10**). For example, for *Aquifex aeolicus* Ribosomal protein L20^63^, the first two axes clearly separate hydrophobic core and surface positions (**SI Figure 11**). The SVD axes therefore correspond to spatially and chemically coherent groups of positions, each associated with its own patterns of tolerated substitutions in the DMS data we investigated.

**Figure 3.**
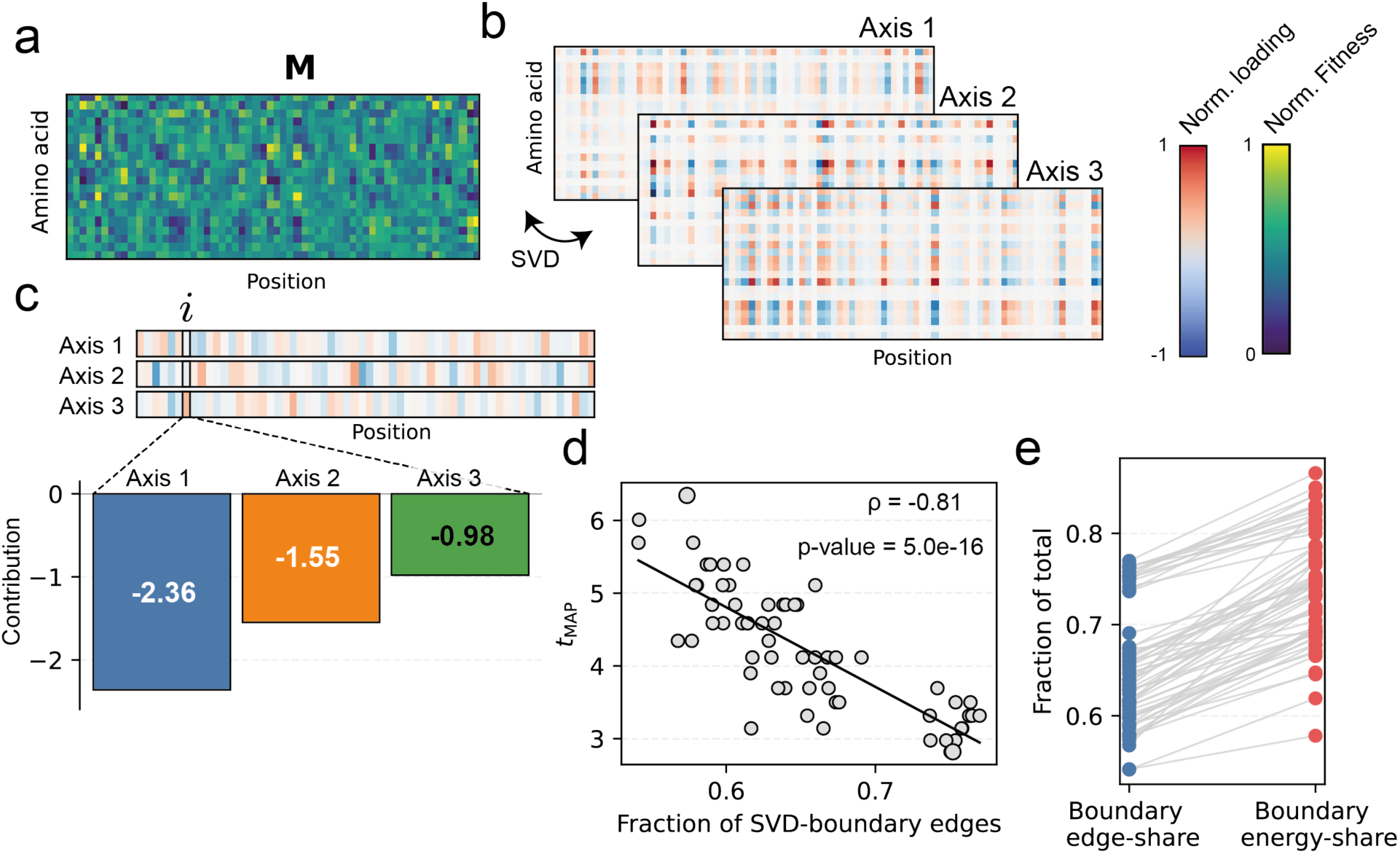
Fitness landscape ruggedness concentrates at SVD-defined substitution regime boundaries. **a**. Position-by-substitution matrix **M** for a representative domain. **b**. Singular value decomposition of M into orthogonal axes that capture independent amino acid-tolerance patterns. **c**. SVD limiter assignment for genotype i: cumulative mutation contributions are summed across axes, and the most negative axis defines the dominant substitution regime. **d**. Domains with a larger fraction of SVD-boundary edges have lower *t*_MAP_. **e**. SVD-boundary edges carry a larger share of Dirichlet energy than expected from their edge share across all 64 domains. For each domain, blue points in **e** show the fraction of tested Hamming-graph edges classified as SVD-boundary edges, and the coupled red points show the fraction of total Dirichlet energy carried by those boundary crossing edges. Energy is enriched along SVD boundary crossing edges, relative to the total share of edges those boundaries account for.

We then asked whether ruggedness in these landscapes concentrates at mutations that switch the dominant constraint. To do this, we first needed to assign each variant in the landscape to one of the SVD axes, identifying which constraint its mutations most violate. For each mutation, we computed a signed contribution to each SVD axis: positive if the substitution is favored by the rule that axis represents, negative if it violates that rule. The magnitude of the contribution scales with how strongly the mutated position participates in the axis. Summing across all a variant’s mutations gives a score for each axis. We assigned the variant to the axis with the most negative summed score, i.e., the constraint its mutations collectively violate most (**Figure 3c; SI Figure 12**). In the sequence graph, edges connect variants that differ by a single substitution. Edges connecting variants assigned to different SVD axes (“boundary edges”) therefore mark mutations that switch the constraint that is assumed most violated according to the SVD. To test this, we asked whether crossing such a boundary involves a substantial change in residue chemistry: boundary edges are strongly enriched for substitutions that change amino acid hydrophobicity or volume (odds-ratio = 1.95, p-value << 0.05), confirming that boundary crossings are biophysically meaningful events and not artefacts of the assignment procedure axis assignment.

Two findings link these boundaries to our global measure of ruggedness, *t*_MAP_. First, the fraction of edges that are boundary edges strongly correlates with *t*_MAP_ across the 64 domains (Spearman’s ρ = -0.81, p-value = 5.0e-16; **Figure 3d**), suggesting that domains in which the dominant constraint changes more frequently across sequence space are more globally rugged. Second, boundary edges carry 1.54-fold more variation than non-boundary edges, when pooled across all domains (p-value = 1.2e-38) (**Figure 3e**). Ruggedness in the DMS stability data is therefore not diffuse; It concentrates at the boundaries between regions of sequence space where fitness is governed by different limiting mutational regimes and patterns. Rugged fitness landscapes are generally those where the limiting mutation regimes change frequently.

### Ruggedness emerges from the composition of individually smooth selective constraints

The 64 DMS stability landscapes showed that ruggedness concentrates on edges where the dominant constraint switches between adjacent sequences, suggesting that the composition of selective constraints, rather than the geometry of the landscape induced by any single one, may be a source of ruggedness. We hypothesized that observable ruggedness can emerge from the geometry of how selective constraints intersect in sequence space, irrespective of how rugged they are in isolation^64^. To formalize this, we represent for each selective constraint *i* a solution set *Si*, defined as the set of sequences whose phenotype under that constraint exceeds a minimum fitness threshold *f*_min_ required to escape purifying selection^65^. For example, *Si* may describe the set of sequences that fold into a stable structure; failure to fold eliminates fitness regardless of how well other constraints are satisfied (**Figure 4a**). Geometrically, the boundary-to-volume ratio of *S_i_* measures how much of *S_i_* sits adjacent to non-viable sequence space: a thin, fragmented solution set has a large boundary relative to its volume, whereas a compact, contiguous one has a small boundary relative to its volume (**Figure 4b, c**). *Si* boundary-crossing edges impose a lower bound on Dirichlet energy when they carry non-zero fitness differences and boundary-rich landscapes are expected to shift fitness variation toward higher-frequency graph modes when those boundaries carry appreciable fitness changes (**SI Text 1**). When m selective constraints act concurrently on the same sequence space, *S_i_* comprises the set of sequences that satisfy all m constraints. Intersecting non-redundant solution sets from m non-redundant constraints can reduce the volume of *S_i_* and introduce additional boundary-crossing edges where different constraints become limiting (**Figure 4d**), providing a route by which composite landscapes can be more rugged than their individual constituents (**Figure 4b, c**). Analogous geometric results have been established in morphological trait space, where competing selection pressures collapse viable phenotypes onto a lower-dimensional Pareto front^66^ and are generally anticipated by Fisher’s Geometric Model (FGM)^58^.

**Figure 4.**
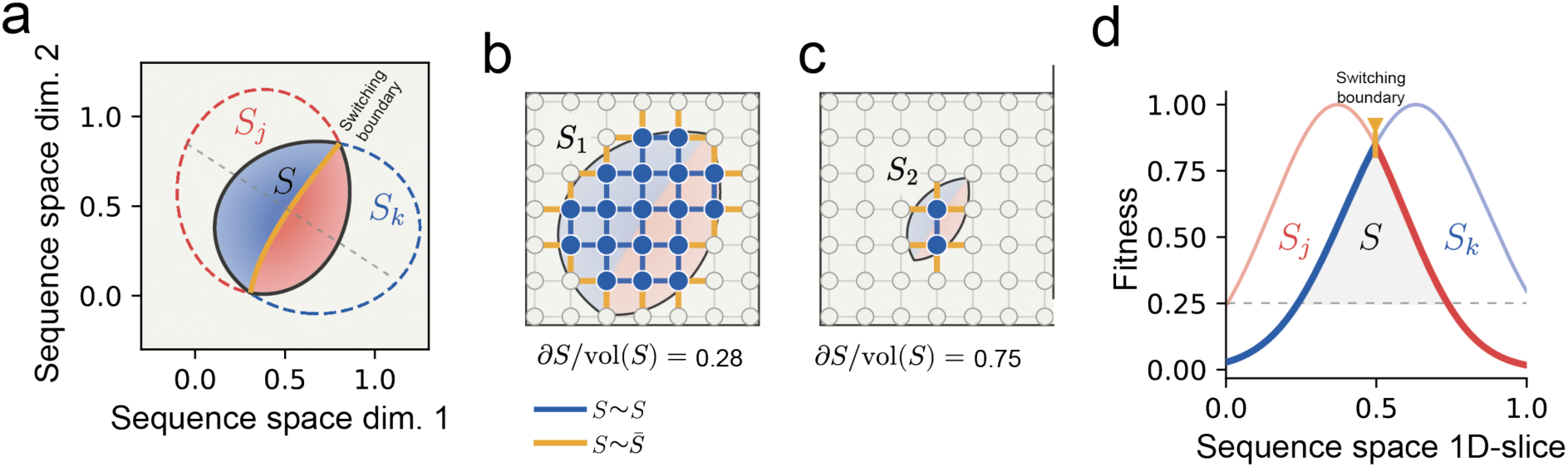
Boundary structure of intersecting selective constraints. **a**. Schematic of two constraint solution sets, *S*_j_ and *S*_k_, whose intersection defines the viable set S. **b, c**. Graph examples showing that smaller or more fragmented solution sets have higher boundary-to-volume ratios, differing by the *f*_min_ on the same example fitness landscape. A larger solution space, as in *S*_1_ produces greater S-to-S adjacency than a smaller solution set, *S*_2_, resulting in a smaller conductance across the S boundary and thus a less spectrally rugged fitness landscape. **d**. One-dimensional fitness slice illustrating how the limiting constraint changes at a switching boundary within S.

We tested this prediction in a minimal model system in which each selective constraint is an additive NK landscape (K=0). Each constituent landscape is therefore entirely free of epistasis and represents a single selective constraint. We assume a hard-constraint regime, in which failure under any one constraint eliminates viability regardless of how well the others are satisfied (a protein that cannot fold gains no fitness from binding its target well), so that the observable fitness of a genotype is the minimum fitness across the constituent landscapes. Under these assumptions, the composite fitness function is the piecewise minimum of the constituent landscapes: at each genotype, the composite fitness is whichever constituent landscape happens to assign the lowest fitness at that point (**SI Figure 13**). We tested whether the piecewise minimum operator is a realistic model of composite selection in a real protein system using a SARS-CoV-2 receptor binding domain (RBD) dataset^67^. Here, a single DMS library of RBD mutants was screened for binding to two antibodies (LyCoV-16, LyCov-555), individually and in combination. Fitness under combined selection was accurately reconstructed from the two single-antibody fitness measurements using the piecewise minimum operator (Pearson’s R^2^ = 0.85). A soft-minimum, which approximates the minimum operation but allows partial compensation between constraints, performed comparably (Pearson’s R^2^ = 0.84), while a multiplicative model, in which fitness contributions from each constraint are multiplied together, performed worse (Pearson’s R^2^ = 0.77; **SI Figure 14**).

We adopted the piecewise minimum for the simulations below on parsimony grounds, since it has no fitted parameters, and confirmed that all qualitative findings are reproduced under soft minimum and multiplicative coupling (**SI Figures 15, 16**). The minimum operator is nonlinear, so composing non-identical additive landscapes necessarily introduces non-additivity into the composite fitness function. We focused on two questions around this non-linearity: how sensitive the emergent ruggedness to the degree of misalignment between constituent constraints is, and where in sequence space is the non-linearity concentrated. We answer these questions using two complementary spectral analyses. First, the graph Fourier transform reveals how fitness variance is distributed across the low- and high-frequency modes of the landscape graph, capturing whether ruggedness is global or local in spectral terms. Second, the per-edge Dirichlet energy reveals where on the graph that variation is spatially concentrated. Both phenomena are captured by the global diffusion scale *t*_MAP_: high-frequency spectral energy reduces *t*_MAP_, and the spatial distribution of Dirichlet energy reveals where on the graph that reduction originates. To control the degree of overlap between constraints, we introduce an alignment parameter α that controls the correlation between the per-site fitness contributions of the constituent landscapes, where α = 0 corresponds to independently sampled landscapes and α = 1 to identical ones (Methods) (**Figure 5a**). Even near-perfect alignment between two constituents (α = 0.90) significantly reduces *t*_MAP_ relative to a single additive landscape (**Figure 5b**), and ruggedness increases monotonically both as constraints become more independent and as more constraints are added (**Figure 5a**). The same effect is visible as epistasis. The Walsh-Hadamard transform decomposes a fitness function into orthogonal components representing additive contributions, pairwise interactions, and progressively higher-order interactions between sites; energy in the higher-order components is the standard spectral signature of epistasis. Composing individually additive landscapes shifts variance into pairwise and higher-order Walsh components, generating the spectral signatures of high-order epistasis even though constituent landscapes are purely additive. This is true even when the constituents are largely redundant (α close to 1; **Figure 5c**), establishing that composition of selective constraints is a general mechanism for the emergence of non-linearity in fitness maps. Filtering out the additive components of coupled NK landscapes (a high-pass filter; **SI Figure 17**) reveals that the non-additive variation in fitness is concentrated on edges that cross constraint boundaries (**Figure 5d, 5e**). Constraint boundaries introduce abrupt fitness gradients between adjacent sequences, generating high-frequency content in the graph Fourier spectrum. Composition of selective constraints can therefore generate ruggedness, even when the constraints are nearly redundant and when each individual constraint is entirely free of specific epistasis. A consistent geometric picture has been reported in environment-dependent fitness landscapes, where ruggedness peaks at intermediate environmental stress because the limiting constraint switches between growth and resistance^68^.

**Figure 5.**
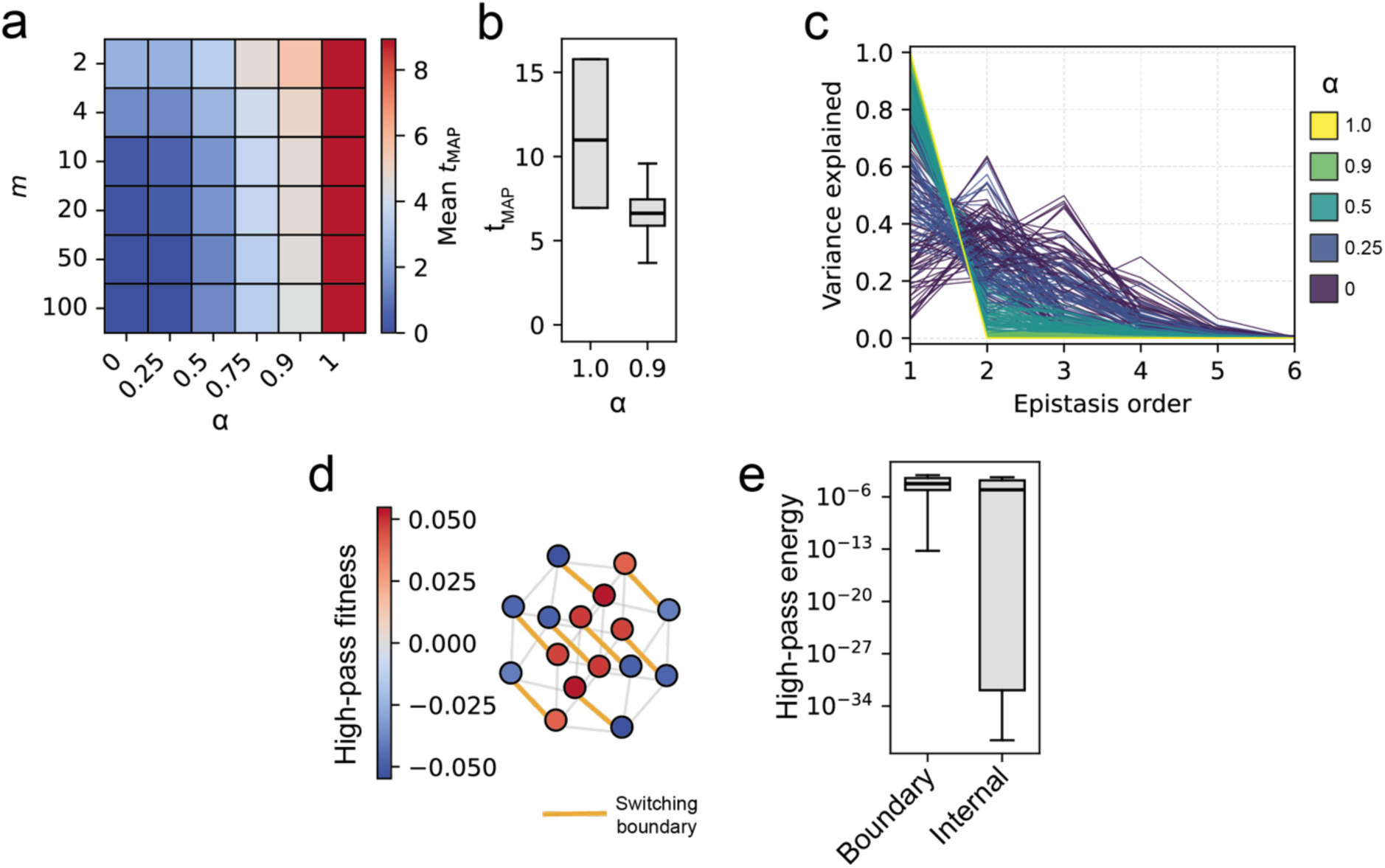
Ruggedness emerges from the composition of individually smooth selective constraints. **a**. Mean *t*_MAP_ for piecewise-minimum composites of additive landscapes across alignment parameter α and number of constraints m. **b**. Near-perfectly aligned constraints (α = 0.90) reduce *t*_MAP_ relative to a single additive landscape. **c**. Walsh-Hadamard decomposition showing pairwise and higher-order variance after coupling additive landscapes. **d**. High-pass fitness on a coupled landscape, with boundary-switching edges highlighted. **e**. The non-additive, filtered signal is enriched on boundary-switching edges relative to internal edges. Emergent ruggedness concentrates locally on the boundary-switching edges under a minimum operator.

### Ruggedness concentrates at regime boundaries in experimental fitness landscapes

We next sought to test whether the emergence of ruggedness from the composition of constraints, and specifically its concentration at constraint boundaries, is observable in experimental fitness landscapes. The DMS unfolding landscapes analyzed above required us to infer the dominant constraint at each position from substitution patterns alone. We therefore turned to datasets in which the latent constraints contributing to fitness are determined directly. Recent applications of the Double Deep Protein-Fragment Complementation Assay (ddPCA) decompose observed binding phenotypes into latent folding and binding components using a fitted thermodynamic model: an SH3 domain with up to 7 combinatorial substitutions^69^ and several KRAS DMS libraries screened against 6 cognate KRAS binding partners^70^. In ddPCA systems, correct folding is assumed necessary but not sufficient for binding, providing a natural hierarchy in which the binding phenotype guarantees folding, but not the converse^71^. We complemented these with the SARS-CoV-2 receptor binding domain (RBD) screened independently against Ly-CoV16 and LyCoV55^67^, which provides a model-independent test in which both constraints are measured directly rather than inferred from a thermodynamic decomposition.

We constructed each dataset as Hamming graphs and annotated each node with both a composite fitness observed under combined selection and the latent fitness values from each individual constraint. For ddPCA datasets, these were the inferred thermodynamic binding and folding contributions^72^; for the RBD datasets they were the per-variant fitness values from the Ly-CoV-16 and Ly-CoV-555 single-antibody DMS experiments (**Figure 6a**). For each node we computed a continuous limiter index that quantifies the imbalance between the two latent constraints. The index is negative when the node is folding-limited, positive when binding-limited, and zero when both constraints are equally limiting (**Figure 6b**). Each edge was then classified by whether the limiter index changes sign across it: regime boundary edges connect nodes whose limiting constraint differs, and intra-regime edges connect nodes that share the same limiting constraint (**SI Figure 18**).

**Figure 6.**
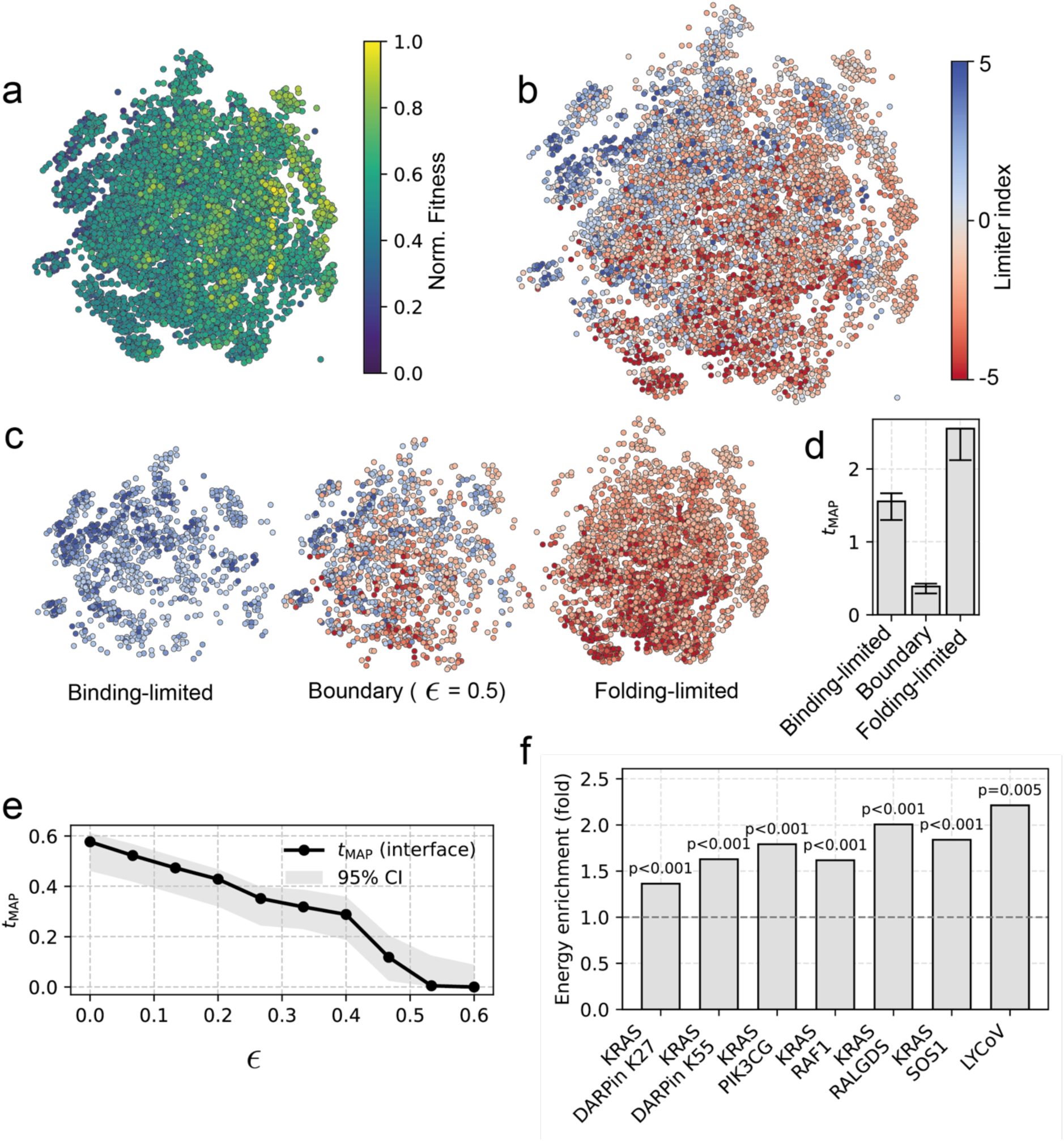
Ruggedness concentrates at regime boundaries in experimental fitness landscapes. **a**. SH3 Hamming graph coloured by normalized composite fitness. **b**. The same graph coloured by limiter index, separating folding-limited, binding-limited, and near-balanced genotypes. **c**. Breadth-first-search subgraphs for binding-limited, boundary, and folding-limited regions. **d**. The SH3 boundary subgraph has a significantly lower *t*_MAP_ than the binding- or folding-limited subgraphs and is thus more rugged. **e**. Boundary *t*_MAP_ decreases as the limiter-index threshold ε is increased; as the boundary requirements become stricter, the boundary subgraph becomes more rugged. **f**. Regime-boundary edges carry excess Dirichlet energy across a range of empirical ddPCA KRAS and explicit two-trait SARS-CoV-2 RBD DMS systems. In experimental fitness landscape, ruggedness is concentrated on mutations that switch the trait that is limiting to selection.

We focused first on the SH3 landscape. The combinatorial and sparse structure of this landscape allowed us to extract similarly sized and connected subgraphs for the regime boundary and for each intra-regime region by breadth-first search (Figure 6c) and to fit *t*_MAP_ independently within each. The regime boundary subgraph was substantially more rugged than either intra-regime subgraph: *t*_MAP_ = 0.38 at the boundary against *t*_MAP_ = 2.6 and 1.6 in the folding-limited and binding-limited regimes respectively (p-values << 0.05) for the pairwise comparisons against the boundary; (**Figure 6d**). Resampling the latent folding and binding values within the uncertainty of the thermodynamic model did not change this result (**SI Figure 19**). The ruggedness at the regime boundary scaled with how sharply the boundary was defined. Restricting the boundary to edges where the difference between latent traits is larger than a constant (ε) reduced the boundary *t*_MAP_ further, thus increasing the observable ruggedness in the partition (**Figure 6e, SI Figure 20**). We also confirmed that the boundary structure is not driven by shared variance between the limiter index and the composite fitness (**SI Figure 21**). Within the binding-limited regime, mutations have approximately balanced effects on the two latent fitness contributions, but transitions into the folding-limited regime occur predominantly through destabilizing substitutions, abruptly steepening the fitness gradient at the boundary (**SI Figure 22**).

We extended this analysis to the KRAS and RBD landscapes. Because these are DMS rather than combinatorial datasets, the graphs could not be cleanly partitioned into connected boundary and intra-regime subgraphs, as for the SH3 domain dataset. We instead used per-edge Dirichlet energy to quantify the fitness variation carried by each edge, and asked what fraction of the total energy in each landscape was accounted for by regime boundary edges. In every system, regime boundary edges contributed a significantly greater share of total Dirichlet energy than expected under a null hypothesis of diffuse ruggedness (p-value < 0.05, 1000 permutations). Normalizing for the proportion of edges classified as regime boundary, these edges carried between 1.63 and 2.25-fold more Dirichlet energy than intra-regime edges (**Figure 6f**). The strongest enrichment was observed in the RBD system, where both constraints are measured directly rather than inferred from a thermodynamic model, indicating that the result does not depend on the ddPCA decomposition. Across the KRAS and RBD landscapes, ruggedness is therefore not uniformly distributed in sequence space; it concentrates at the edges where the limiting constraint differs between the two connected nodes.

## Discussion

Our results show that rugged protein fitness landscapes can arise from constraints that are individually simple. A protein under selection must satisfy several biophysical requirements at once, such as folding stability and binding affinity, and the set of sequences that satisfies all of them is the intersection of the sets that satisfy each individually. Ruggedness concentrates at the edges where the most limiting constraint switches. More generally, the idea that multiple underlying constraints can generate epistasis or landscape complexity is not new: related intuitions appear in FGM and in prior work on thermodynamic and functional epistasis^58,68,73–76^. Our results, however, provide insight on a specific mechanism through which this can occur: the concentration of high-frequency fitness signals into the boundary-switching edges where the most limiting functional constraint changes, which we observe across empirical fitness landscapes. A broader implication is that ruggedness need not be globally diffuse across a fitness landscape but can instead be locally concentrated in regions where the effective limiting constraint changes over sequence space, revealing the hidden biophysical constraints on fitness.

An important consequence of our results is that protein folding landscapes should be systematically smoother than the landscapes for any function that depends on folding. Correct folding is necessary but not sufficient for many biological functions — a catalytically competent protein is by necessity folded, but a folded protein is not by necessity catalytically competent^53,77,78^. The folding landscape is therefore a superspace of any functional landscape that depends on folding, and the set of sequences that will fold is at least as large as the set that can facilitate a function requiring correct folding. More precisely, our results anticipate that the lower bound on ruggedness of folding stability is less than or equal to that of any function that depends on folding, consistent with recent observations that folding stability has a relatively simple genetic architecture^16,17^.

The localisation of ruggedness at regime boundaries has consequences for protein engineering and for natural evolution. Because fitness changes abruptly over short mutational distances at these boundaries, rugged fitness landscapes are difficult to learn, and predictive models fail to generalize across edges where the limiting constraint changes^34,79^. If training data is drawn predominantly from within a single regime, held-out performance will look better than it should, because the boundary is under-sampled. Spectral cuts of the fitness graph provide a way to locate regime boundaries directly from the data, which suggests two engineering strategies: fit separate models within each regime and combine them at the boundaries, or weight the training procedure so that boundary edges are not learned in the same way as intra-regime edges. Indeed, there has been considerable work on adapting deep learning protein design models to learn the signals of underlying spectral epistasis and fitness landscape ruggedness from experimental data^34,80–82^. Whether spectral graph methods can be used to identify such boundaries in practice, and whether this improves protein design outcomes, will be the focus of our future work.

There are several limitations of our study that should be noted. In experimental systems, boundary edges account for a significant yet modest (1.63 to 2.25-fold) enrichment in Dirichlet energy relative to intra-regime edges. Thus, other mechanisms contribute to the residual variation, including direct interactions between specific residues, higher-order structural effects, and the consequences of proteins existing as a distribution of conformations rather than a single structure^7,8,73–75^. What we describe is therefore only a part of the complexity in the sequence-fitness relationship, and not its absolute origin. Additionally, the boundary analysis in ddPCA systems depends on latent folding and binding contributions inferred from a thermodynamic model^69,72^; while resampling within model uncertainty did not quantitatively change our results or their interpretation, the boundary identification inherits whatever biases exist in that model, including the reasonable assumption that folding is a necessary condition for binding. Relatedly, the SVD-defined constraint axes used in the DMS stability analysis are scale-dependent: they group positions that covary in substitution tolerance at the granularity of a single domain and would partition differently if applied to landscapes spanning multiple folds, functions or mutational scales. The SARS-CoV-2 RBD landscape^67^, in which both constraints are measured directly rather than inferred, gives a result that is consistent with those inferred from a thermodynamic model, indicating that our findings are not artefactual; however, the rarity of published experimental systems where multiple traits are assessed both in isolation and in concert make generalisations to diverse systems difficult to establish. Subject to these caveats, the central finding of this work is that the composition of individually simple selective requirements is sufficient to generate localized ruggedness, with a predictable spatial signature at the regime boundaries where the limiting constraint differs between adjacent sequences.

## Methods

### Fitness Landscape Hamming Network Graphs

All landscapes were modelled as undirected NetworkX (v3.1) graphs G = (V, E), with one node per unique sequence. Hamming graphs were constructed such that two nodes *i* and j are connected only when their sequences differed by a single substitution. Hamming adjacency for binary sequence spaces was computed using a bitwise XOR lookup, whereas multiallelic sequence spaces were constructed using a masked radix-encoding scheme that identifies all Hamming-distance-1 neighbours. This construction was used for NK landscapes, complete substitution datasets, megascale folding datasets, and the SH3 core landscape.

### NK landscape construction

NK landscapes were generated using the standard NK model with N binary variable sites and K epistatic neighbours per site. For each site *i*, a random fitness contribution table was constructed mapping the 2^K+1^ possible allelic states of site *i* and its K neighbours to fitness contributions drawn from a uniform distribution centred at zero. The fitness of a genotype was computed as the normalised sum of all per-site contributions: 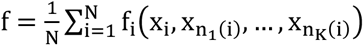. Epistatic neighbours were assigned randomly unless a custom adjacency matrix was provided.

### Sequence Evolution Simulations and kNN Graphs

Phylogenetic simulations were conducted using AliSim^49^, as implemented in IQ-TREE3^83^. Alignments were simulated over random phylogenetic topologies, using the LG^84^ replacement matrix with a gamma rate heterogeneity parameter with 4 rate categories. Network graphs were constructed from these simulated alignments as kNN graphs, with adjacency computed using the SciPy (v1.11.4) BallTree algorithm^85^.

### Elementary Landscapes

Elementary landscapes were first constructed as *kNN* or Hamming graphs. For the *j^th^* elementary landscape, the signal at the *i^th^* node was assigned as the *i^th^* element of the *j^th^* eigenvector of the unnormalized graph Laplacian *L*. Sparse spectral decomposition of *L* up to the *j^th^* eigenpair was performed in SciPy using the eigsh algorithm.

### Gaussian Markov Random Field

We modelled covariation in the fitness landscape as multivariate Gaussian Markov random fields (GMRFs). GMRFs are spatial statistical models that can be described by a graph; the fitness value of all nodes is jointly distributed as a Gaussian distribution, and edges describe the conditional dependencies between nodes. Conditional dependencies in the GMRF are captured by the sparse precision matrix *Q*, where *i*, *j^th^* entry is 0 if nodes *x_i_* and *x_j_* are conditionally independent. The covariance matrix of the GMRF, Σ, is the inverse of the precision matrix, *Q*^-1^, defining the GMRF as:

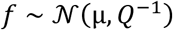

Where µ is the average signal.

### Heat Diffusion Kernel

The heat diffusion kernel^38^ was computed in the spectral domain of the normalized Graph Laplacian. The normalized graph Laplacian, *L̂*, was computed as:

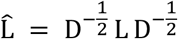

Where *L* is the unnormalized Laplacian, *L* = *D* − *A*, *D* is the diagonal degree matrix and *A* is the adjacency matrix, weighted by element wise inverse of the Hamming distance matrix. The raw heat kernel value, ℎ*_i_* was then computed as:

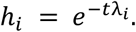

Where λ*_i_* are the eigenvalues of *L̂*, and *t* is the heat diffusion scale. Raw heat kernel values, ℎ*_i_* were scaled to match the observed variance in the fitness vector, *f*. Specifically, let *f* be the observed fitness vector and *L̂* be its projection onto the Laplacian eigenbasis. We compute the following scaling factor:

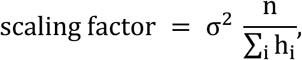

and set

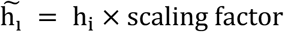

Here, σ^2^ is the observed variance in *f*, and n is the dimension of *f*. The covariance matrix in the spectral domain is diagonal, with entries 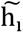. The log likelihood of an observed signal being sampled under the GMRF parameterized with diffusion scale *t* then evaluates to:

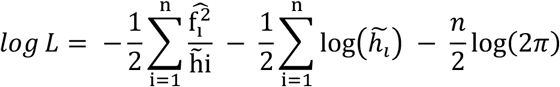

Where 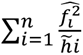 is the quadratic form, 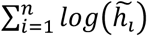 is the log determinant and 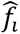 are the coefficients of f in the in the basis of the eigenvectors of *L̂*.

Transformation of the heat diffusion kernel to the spectral domain of the graph Laplacian gives us

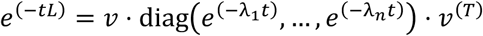

where *v* and λ*_i_* are eigenvalues and eigenvectors of the Laplacian *L̂*. This spectral form shows how the kernel attenuates high frequency eigenmodes of the graph Laplacian by dampening the rugged components of the fitness in the domain of the graph once transformed back into the sequence domain.

### Bayesian fitting of heat diffusion scale parameter

The posterior over *t* was approximated on a logarithmically spaced grid of 512 values between *t*_min_ = 10^-10^ and *t*_max_ = 10^2^, using a uniform prior on this interval. Grid points were weighted by their spacing (trapezoidal quadrature) before normalisation to approximate the continuous posterior density. The reported *t*_MAP_ is the grid point with maximal posterior density. The accompanying 95% credible interval was obtained by interpolating the 2.5% and 97.5% quantiles of the cumulative posterior distribution over the grid.

### Spectral bipartitioning of fitness landscapes

To test whether fitness landscape ruggedness is diffusely distributed or can be localised to a small number of edges, we applied a fitness-weighted spectral bipartitioning to each DMS Hamming graph. For each edge (*i*, *j*), the per-edge Dirichlet energy *E_ij_* = (*f_i_* – *f_j_*)^2^ was computed. Edges were then assigned a weight inversely proportional to their fitness gradient:

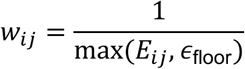

where *ε*_floor_is the 5th percentile of positive edge energies, floored at 10^-12^, to prevent division by zero or extreme weights from near-zero-gradient edges. This weighting scheme ensures that the Fiedler vector of the resulting weighted Laplacian preferentially cuts through high-energy (large fitness gradient) edges, because edges with large fitness differences receive small weights and are therefore cheap for the spectral partition to sever.

The Fiedler vector (the eigenvector corresponding to the second-smallest eigenvalue of the weighted graph Laplacian) was computed using the tracemin_pcg algorithm as implemented in NetworkX. Nodes were assigned to two partitions by thresholding the Fiedler vector at its median value, with a minimum partition size of 2 nodes enforced. Edges connecting nodes in different partitions define the spectral cutset. We note that this bipartitioning depends on the fitness signal through the edge weighting and is therefore not a purely topological partition.

The Dirichlet energy enrichment of the spectral cutset was computed as the ratio of the fraction of total Dirichlet energy on cutset edges to the fraction of total edges in the cutset:

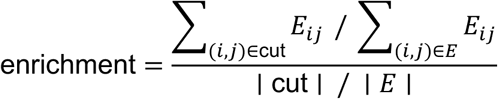

An enrichment greater than 1 indicates that cutset edges carry a disproportionate share of total fitness variation. The statistical significance of the enrichment across domains was assessed by a one-sided Wilcoxon signed-rank test against a null hypothesis of enrichment = 1.

### Shell distance profiles from the spectral cut

To assess whether Dirichlet energy decays with distance from the spectral cutset, we computed shell distance profiles for each domain. After removing the cutset edges from the graph, the shortest-path distance from each remaining node to the nearest node incident on a cutset edge was computed using multi-source Dijkstra’s algorithm (unweighted). Each non-cutset edge was then assigned a shell distance equal to 1 + min(*d_u_*, *d_v_*), where *d_u_* and *d_v_* are the distances of its endpoints to the nearest cutset node. Cutset edges were assigned shell distance 0. Per-edge Dirichlet energy was then aggregated by shell distance to produce a profile of mean edge energy as a function of distance from the spectral cut.

### SVD of the substitution-effect matrix, M

For each of the 64 DMS datasets, a position-by-amino acid substitution matrix **M** was constructed, where rows correspond to positions in the protein sequence and columns correspond to the 20 canonical amino acids. Each entry M_p,a_ records the mean fitness effect of substituting amino acid *a* at position *p*, as measured from single-mutant DMS scores. Wild-type (self-substitution) entries were excluded by setting them to missing. Rows with fewer than 5 measured substitutions were discarded. Missing values were imputed with the row mean, and each row was centered by subtracting its mean.

The singular value decomposition (SVD) of the cantered matrix was computed as M = USV^T^ using ‘numpy.linalg.svd’. The number of retained components *k* was determined as the minimum number of singular values required to explain at least 90% of the total variance (sum of squared singular values), subject to a minimum of 2 and a maximum of 5 components. The row scores were computed as U_;;k_ ⋅ diag(S_:k_), giving a position-by-component matrix where each entry reflects the contribution of each SVD axis at each position.

### SVD limiter boundary assignment

Each genotype in a DMS Hamming graph was assigned to a dominant SVD component (substitution regime) as follows. For each mutation in a genotype relative to the wild-type sequence, the contribution to each SVD component *k* was computed as the product of the row score at that position and the corresponding entry of the right singular vector:

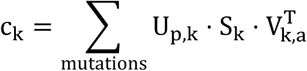

where *p* is the mutated position and *a* is the substituted amino acid. Each genotype was assigned to the SVD component with the minimum (most negative) cumulative contribution, representing the axis under which the genotype’s mutations score most poorly. Genotypes where the two lowest-scoring components differed by less than 10^-8^ were labelled ambiguous and excluded from boundary analysis. Wild-type and genotypes with no covered mutations were similarly excluded.

An edge on the Hamming graph was classified as an SVD boundary edge if its two endpoint genotypes were assigned to different SVD components. The fraction of total edges classified as SVD boundary edges was computed for each domain.

### Coupled NK landscape construction

Coupled fitness landscapes were constructed by composing *m* constituent NK landscapes under a pointwise minimum operator. Each constituent landscape was generated as an NK landscape with specified N and K parameters. To control the degree of correlation between constituent landscapes, an alignment parameter α ∈ [0,1] was used. A single shared NK landscape was first generated. Each of the *m* constituent signals were then constructed as a convex combination of the shared signal and an independently generated NK signal:

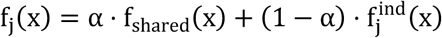

Where f_shared_ is the shared landscape and 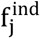 is an independent landscape. When *α* = 1, all constituents are identical (no ruggedness from composition); when *α* = 0, constituents are independently generated. The observable fitness of each genotype in the coupled landscape was defined as the pointwise minimum over all constituent signals:

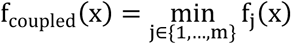

The boundary assignment for each genotype was determined by argmin *_j_f_j_*(*x*), identifying which constituent constraint is most limiting. This construction models multi-constraint selection in which the fitness of a genotype is determined by its most limiting trait.

### Walsh-Hadamard analysis of epistasis

For binary sequence landscapes on full Hamming cubes 2^N^, epistasis was quantified using the Walsh-Hadamard transform (WHT). The fitness vector was reordered to lexicographic genotype order and transformed using the fast Walsh-Hadamard transform (FWHT), normalised by 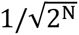. Walsh coefficients were grouped by interaction order (the number of sites involved in each epistatic term, determined by the popcount of the corresponding binary index), and the variance explained by each order was computed as the sum of squared coefficients at that order.

### Graph Fourier filtering of coupled landscapes

To isolate the non-additive (epistatic) component of a coupled fitness landscape, the fitness signal was projected onto the eigenbasis of the graph Laplacian via the graph Fourier transform. For a binary Hamming graph with N variable sites, the first N(A − 1) + 1 eigenmodes (where A = 2 is the alphabet size) span the additive subspace (the constant mode plus N single-site modes). A spectral filter was constructed by setting coefficients of these low-frequency modes to zero and retaining all higher-order modes. The filtered signal was reconstructed by inverse graph Fourier transform, yielding the high-pass (non-additive) component of fitness. Per-edge Dirichlet energy was then computed on this residual signal to assess the spatial distribution of non-additive fitness variation, specifically whether it concentrates at boundary-switching edges.

### Per-edge Dirichlet energy

The Dirichlet energy of a fitness signal f on a graph G = (V, E) quantifies the total squared variation of f along the edges of the graph. For each edge (i, j) ∈ E, the per-edge Dirichlet energy is defined as:

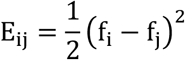

The total (global) Dirichlet energy is the sum over all edges, normalised by the number of nodes:

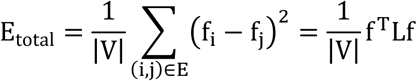

Where *L* is the unnormalized graph Laplacian. Per-edge Dirichlet energy was used throughout to quantify the local contribution of individual edges to global fitness variation.

### Limiter index construction

For ddPCA datasets, a continuous limiter index was constructed for each genotype to quantify whether its composite fitness is primarily constrained by folding stability or binding affinity. Latent folding Δ*G_f_* and binding Δ*G_b_* contributions were obtained from the fitted thermodynamic model. The limiter index was defined such that negative values indicate folding-limited genotypes (folding contribution more negative than binding), positive values indicate binding-limited genotypes, and values near zero indicate genotypes approximately equally limited by both traits. Edges on the Hamming graph where the limiter index changes sign between neighbouring genotypes were classified as boundary-crossing edges.

For the SARS-CoV-2 RBD dataset, the limiter index was constructed analogously using the two directly measured fitness values from individual Ly-CoV-16 and Ly-CoV-555 DMS experiments, rather than latent thermodynamic contributions. This provides a model-agnostic test of boundary-localised ruggedness, as both constituent constraints are experimentally characterised rather than inferred from a thermodynamic decomposition.

### SH3 subgraph partitioning

The sparse combinatorial SH3 landscape was partitioned into boundary and intra-regime subgraphs by breadth-first search (BFS). Nodes were first classified as boundary-adjacent (incident on at least one boundary-crossing edge) or intra-regime (all incident edges within a single limiter regime). Connected components of boundary-adjacent and intra-regime nodes were identified by BFS, yielding similarly sized and connected subgraphs. The diffusion scale t_MAP_ was then computed separately for each subgraph.

## Data and Code Availability

All code and data necessary to reproduce results presented throughout the main and supplementary texts are available at the GitHub: https://github.com/RSCJacksonLab/composition-of-constraints and Zenodo: 10.5281/zenodo.20882872

## Acknowledgements

We acknowledge the ARC Centre of Excellence for Innovations in Peptide and Protein Science (CE200100012) the ARC Centre of Excellence in Synthetic Biology (CE200100029). This research was undertaken with the assistance of resources from the National Computational Infrastructure (NCI Australia), an NCRIS enabled capability supported by the Australian Government.

