## Supporting Information for "The composition of biophysical constraints generates complex, rugged regions of protein fitness landscapes"

Spence M. A.[1,2,3,4]*, Sandhu M.[1], Matthews D. S.[1,2], Nichols J.[3,4], Stone E.[3,4], Jackson C. J.[1,2,3,5]*

1. Research School of Chemistry, Australian National University, Canberra, Australian Capital Territory 2601, Australia
2. ARC Centre for Innovations in Peptide and Protein Science, Australian National University, Canberra, Australian Capital Territory 2601, Australia
3. Research School of Biology, Australian National University, Canberra, Australian Capital Territory 2601, Australia
4. ARC Centre of Excellence in Mathematical Analysis of Cellular Systems, Australian National University, Canberra, Australian Capital Territory, 2601, Australia
5. ARC Centre of Excellence in Synthetic Biology, Australian National University, Canberra, Australian Capital Territory 2601, Australia

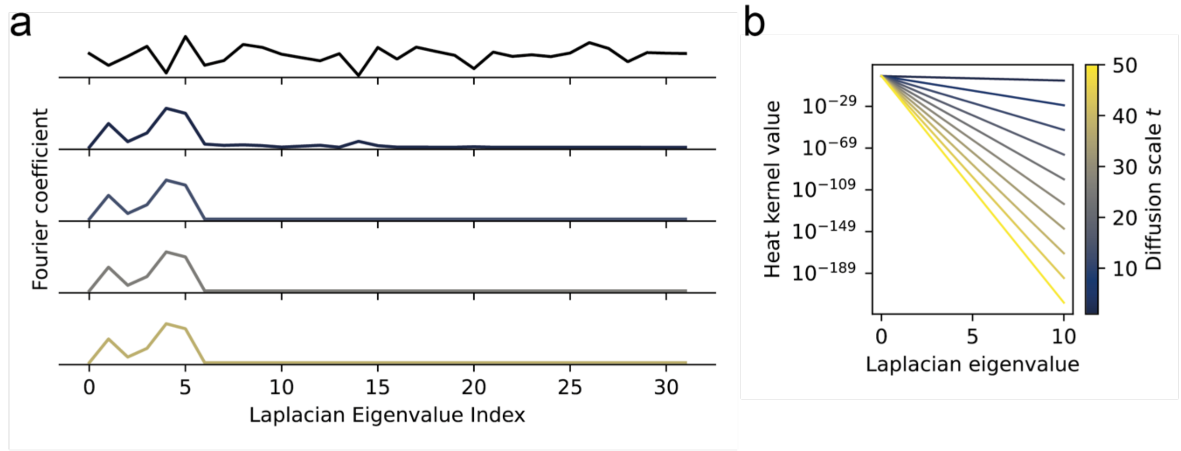


**SI Figure 1.** Heat-kernel attenuation of high-frequency Laplacian eigenmodes. a. Example graph-Fourier coefficients before and after smoothing at increasing diffusion scales, showing preferential loss of high-index modes. b. Heat-kernel weights decrease more steeply with Laplacian eigenvalue as t increases.

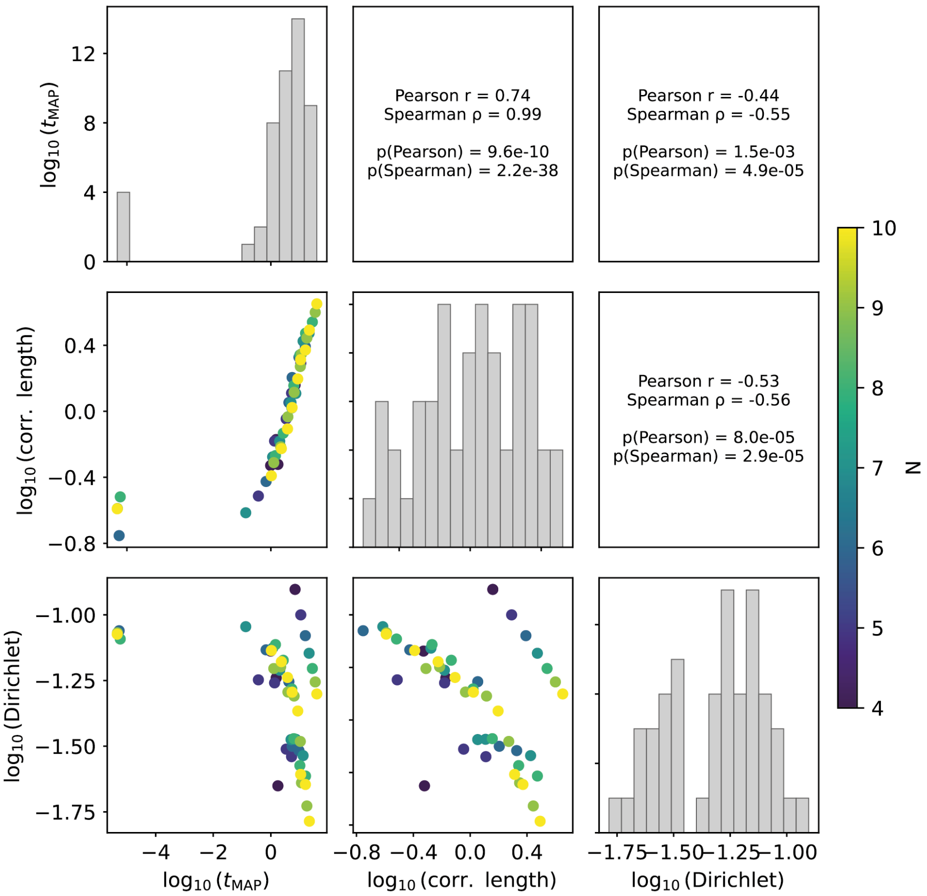


**SI Figure 2.** Concordance among Laplacian-based ruggedness indicators on dense NK landscapes. Pairwise correlogram comparing $t_{MAP}$, random-walk correlation length, and Dirichlet energy; diagonal panels show marginal distributions and off-diagonal panels show pairwise associations coloured by N.

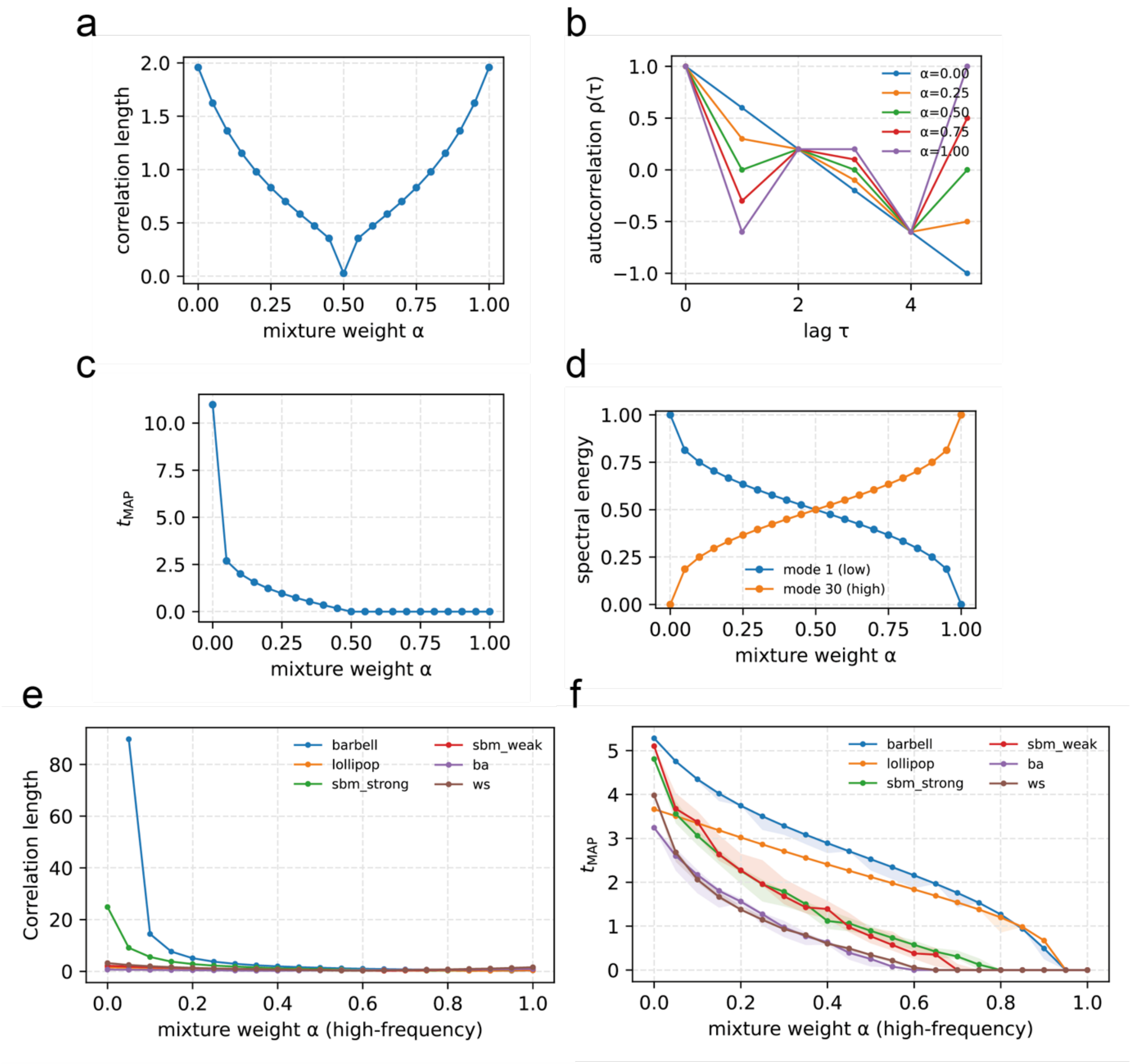


**SI Figure 3.** Laplacian-based ruggedness indicators on spectrally imbalanced model landscapes. a-d. Two-mode landscape showing correlation length, random-walk autocorrelation, $t_{MAP}$, and spectral energy as the mixture weight between low- and high-frequency eigenmodes changes. e, f. Comparison of correlation length and $t_{MAP}$ across sparse graph topologies as high-frequency mixture weight increases.

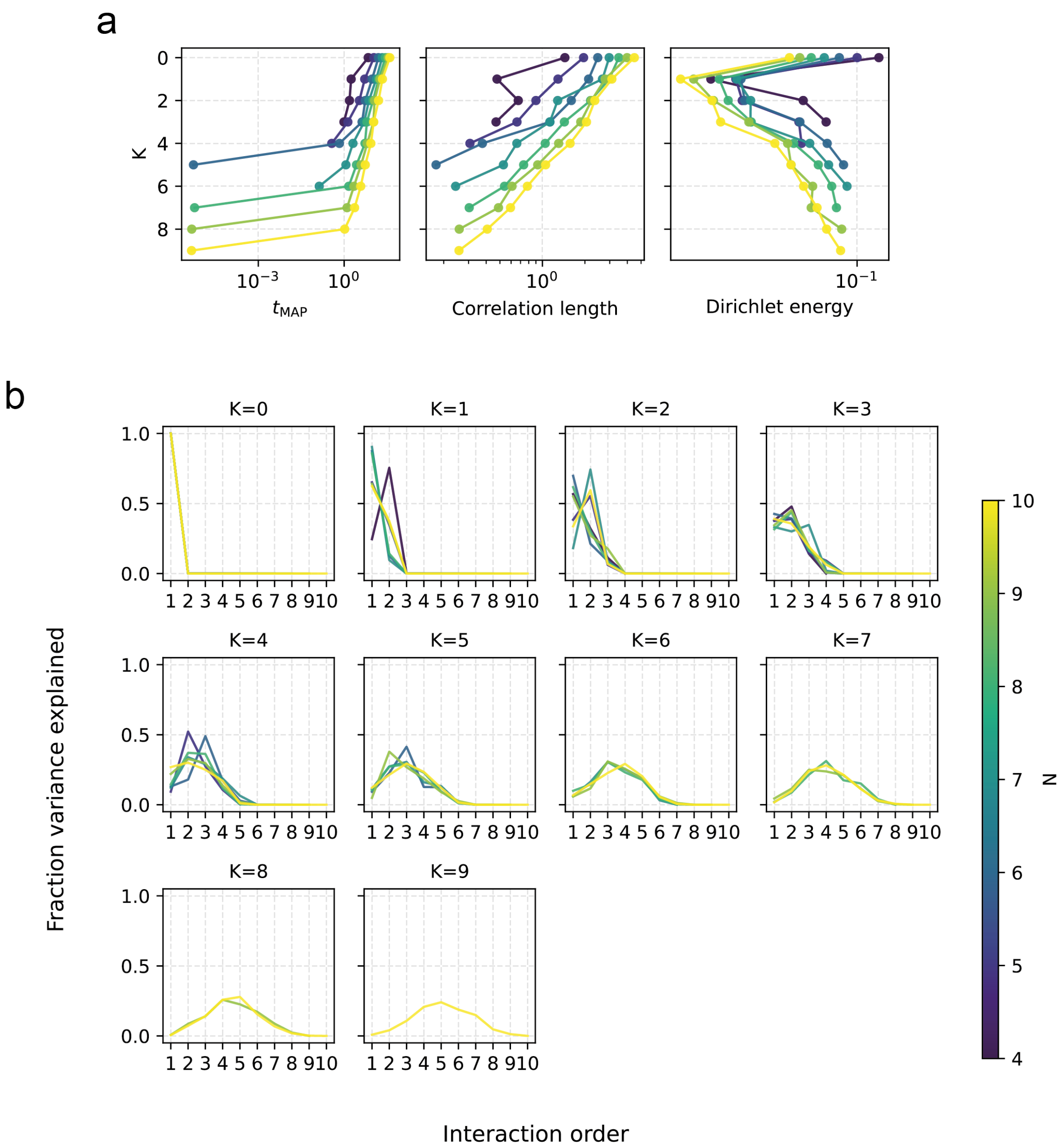


**SI Figure 4.** Epistasis and geometric ruggedness across dense NK landscapes. a. $t_{MAP}$, random-walk correlation length, and Dirichlet energy as functions of K across sequence-space sizes N. b. Walsh-Hadamard variance by interaction order for each K and N.

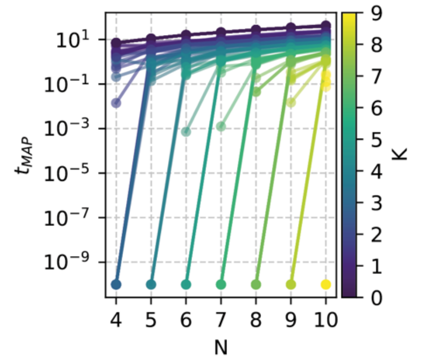


**SI Figure 5.** Dependence of $t_{MAP}$ on sequence-space size and epistasis in dense NK landscapes. $t_{MAP}$ is plotted across N for each NK interaction parameter K, including the size dependence observed when K = 0.

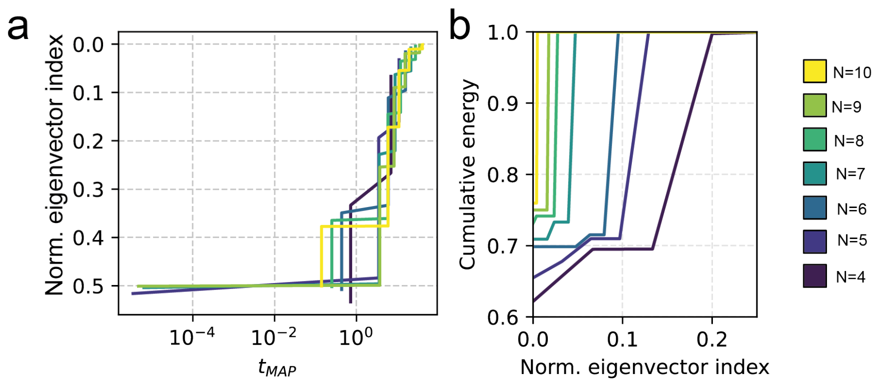


**SI Figure 6.** Normalized spectral position accounts for the apparent N dependence of NK ruggedness. a. $t_{MAP}$ plotted against normalized Laplacian eigenvector index across N. b. Cumulative spectral energy as a function of normalized eigenvector index.

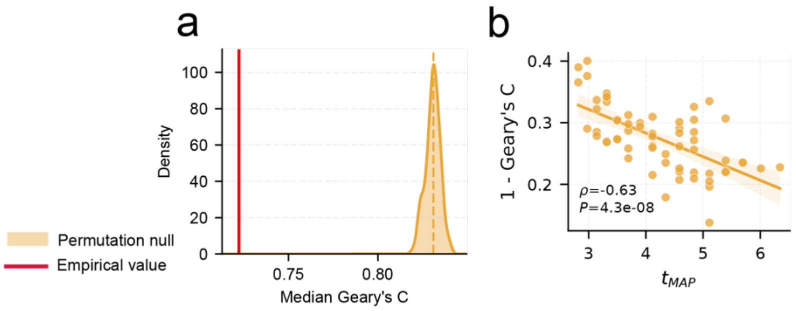


**SI Figure 7.** Spatial autocorrelation of high-energy edges in DMS stability landscapes. a. Node-permutation null distribution for median Geary's C, with the empirical value marked in red. b. Edge-energy autocorrelation, measured as 1 - Geary's C, decreases with $t_{MAP}$ across domains.

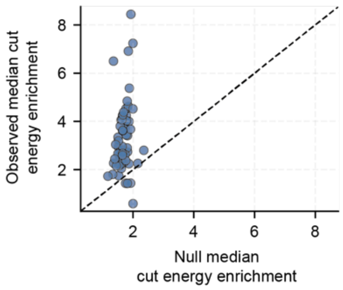


**SI Figure 8.** Cut-energy enrichment depends on the fitness signal rather than topology alone. Observed median spectral cut enrichment is compared with the median enrichment after node-permutation nulls for each DMS domain; points above the diagonal have greater empirical enrichment than expected from topology.

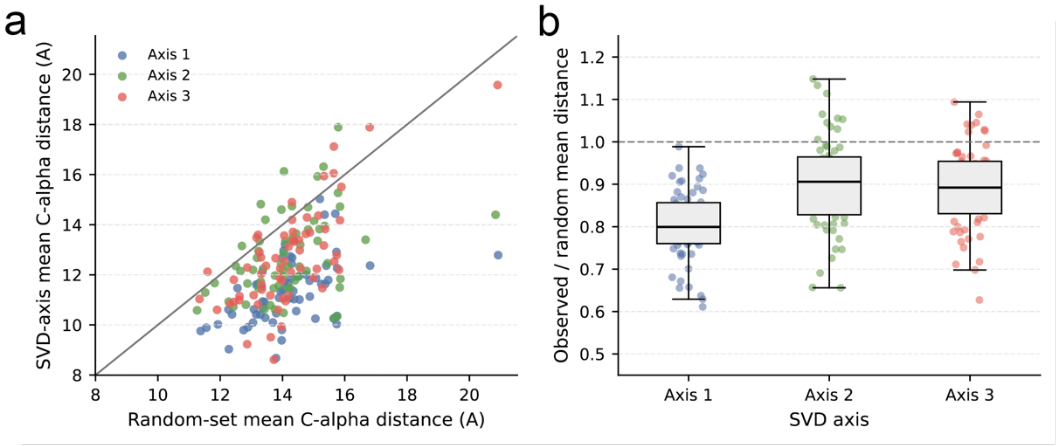


**SI Figure 9.** SVD axes define spatially coherent groups of positions. a. Mean C-alpha distances among positions assigned to each SVD axis compared with random position sets. b. Observed-to-random mean distance ratios for axes 1-3, with values below one indicating spatial clustering.

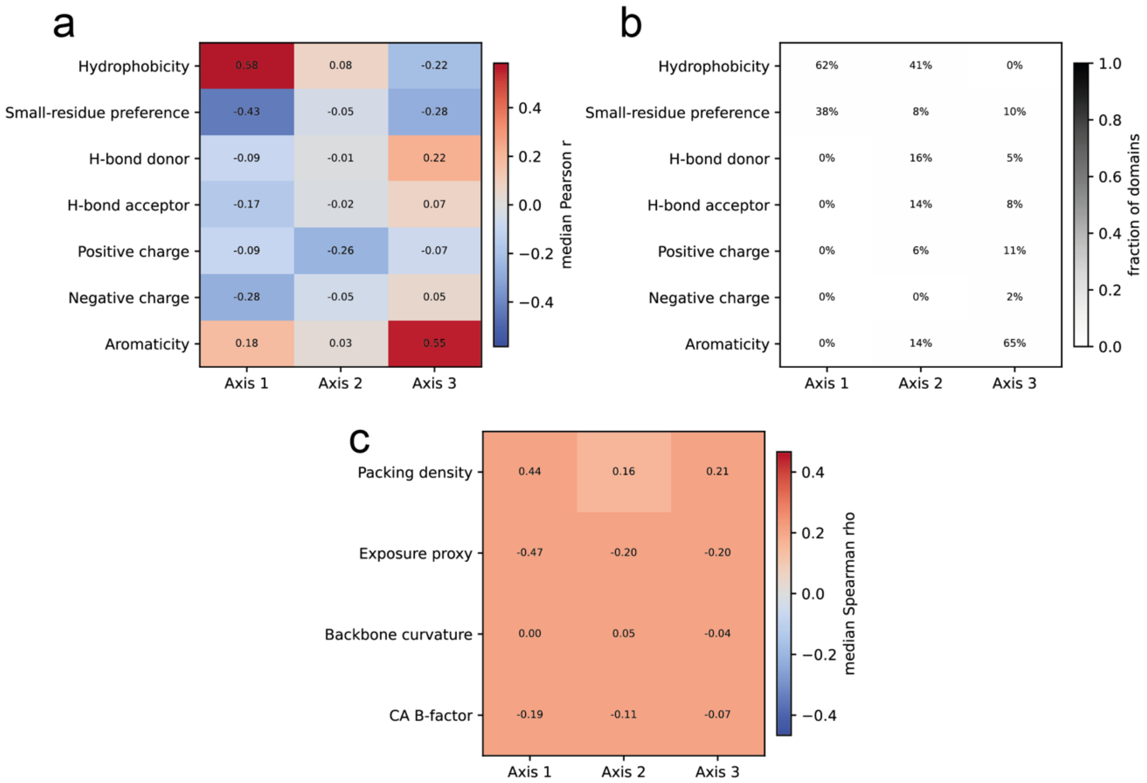


**SI Figure 10.** Chemical and structural correlates of SVD axes across DMS domains. a. Median correlations between SVD axes and amino-acid properties. b. Fraction of domains in which each amino-acid property is associated with each axis. c. Median correlations between SVD axes and structural features of positions.

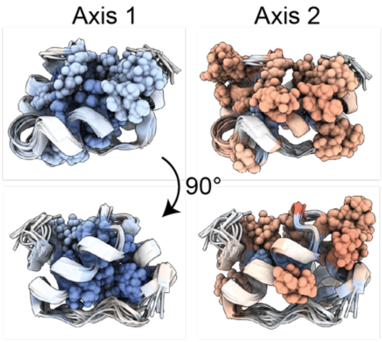


**SI Figure 11.** Structural localization of SVD axes in a representative domain. Axis-1 and axis-2 position loadings are mapped onto the protein structure and shown in two orientations separated by 90 degrees.

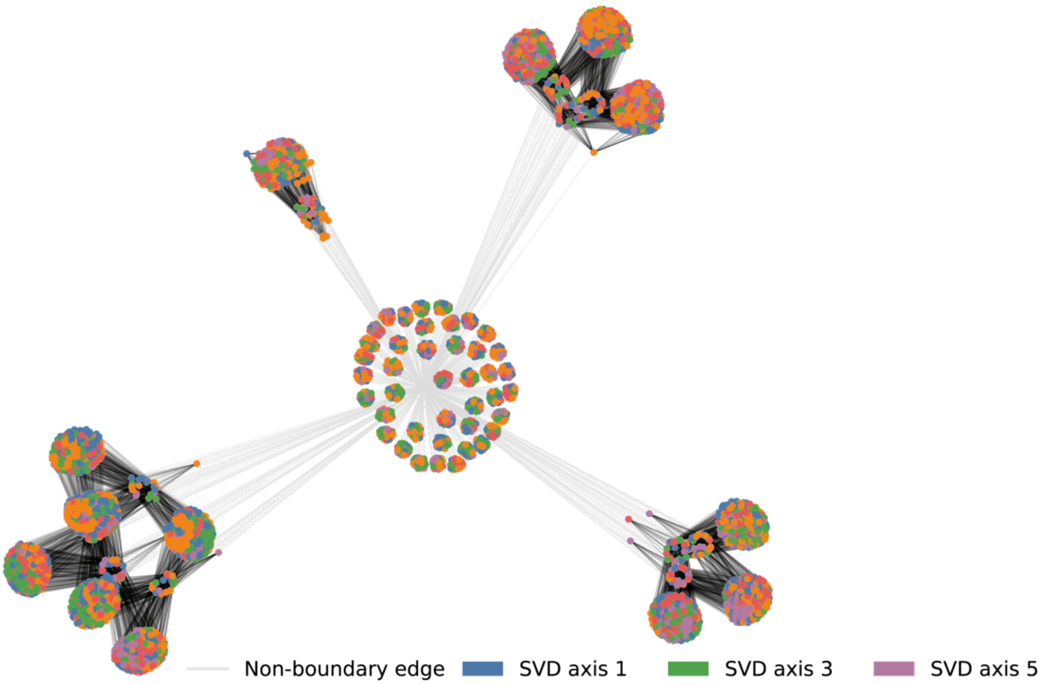


**SI Figure 12.** Example SVD-regime assignment on a domain Hamming graph. Nodes are coloured by assigned SVD axis; black edges connect genotypes assigned to different axes and therefore mark SVD-boundary edges.

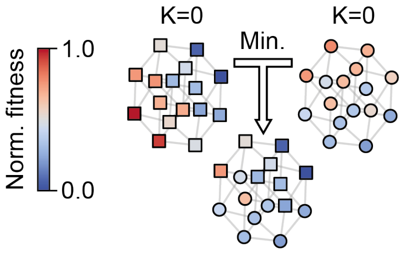


**SI Figure 13.** Piecewise-minimum composition of two additive NK landscapes. Two K = 0 constituent landscapes are combined by taking the lower normalized fitness at each genotype, producing a composite landscape whose limiting constraint can switch across sequence space.

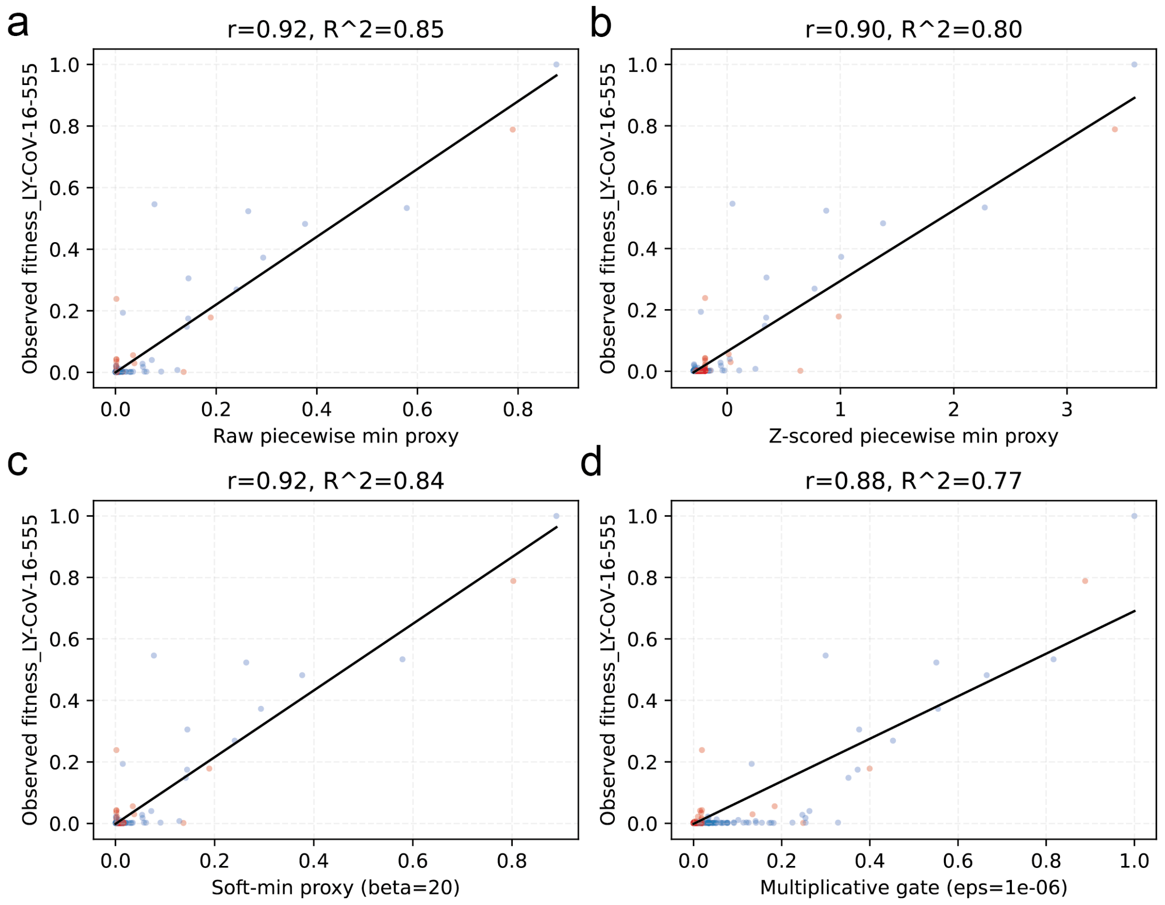


**SI Figure 14.** Predicting combined antibody-selection fitness from single-constraint measurements. Observed Ly-CoV-16/555 fitness is compared with a. raw piecewise-minimum, b. z-scored piecewise-minimum, c. soft-minimum, and d. multiplicative-gate proxies built from the single-antibody landscapes.

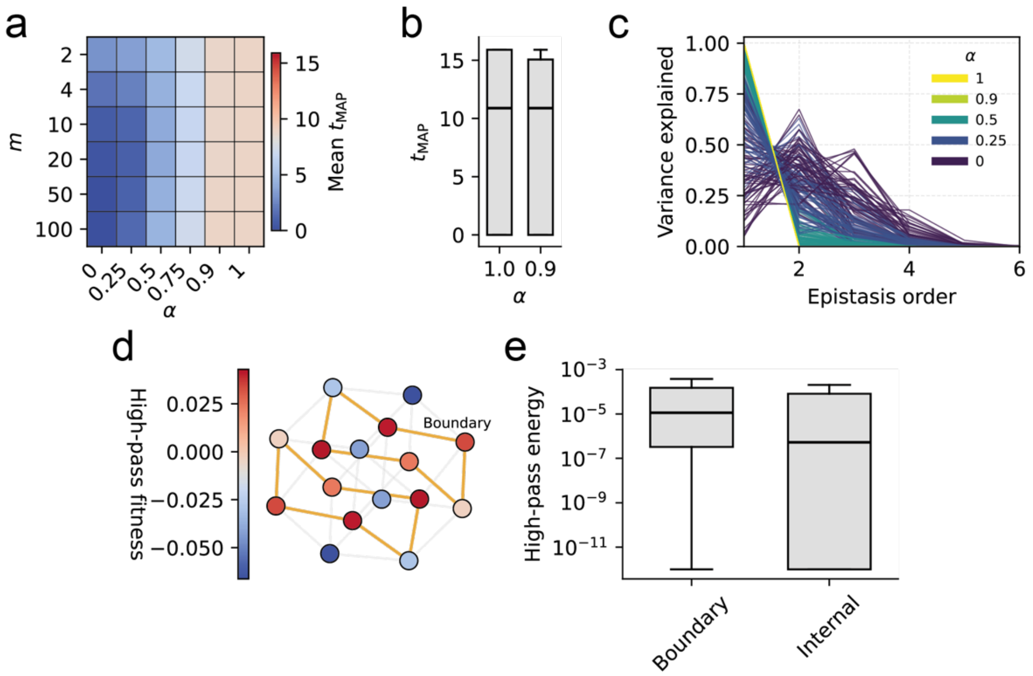


**SI Figure 15.** Soft-minimum coupling reproduces the composite-constraint ruggedness pattern. a. Mean $t_{MAP}$ across alignment α and number of constraints m. b. $t_{MAP}$ for α = 1.0 and α = 0.9. c. Walsh-Hadamard variance by epistasis order. d. High-pass fitness on a coupled landscape with boundary edges highlighted. e. High-pass energy on boundary versus internal edges.

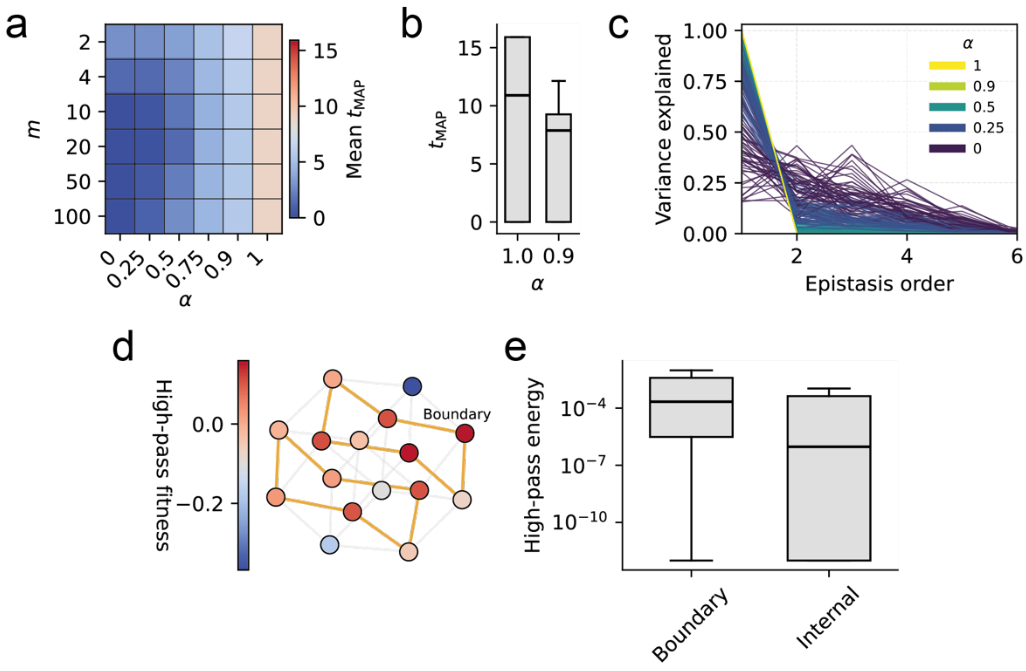


**SI Figure 16.** Multiplicative coupling reproduces the composite-constraint ruggedness pattern. a. Mean $t_{MAP}$ across alignment α and number of constraints m. b. $t_{MAP}$ for α = 1.0 and α = 0.9. c. Walsh-Hadamard variance by epistasis order. d. High-pass fitness on a coupled landscape with boundary edges highlighted. e. High-pass energy on boundary versus internal edges.

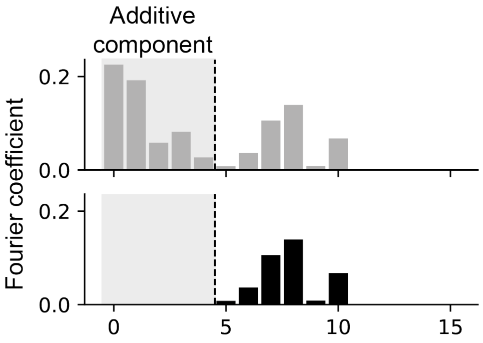


**SI Figure 17.** High-pass filtering removes additive components from composite landscapes. Graph-Fourier coefficients before and after filtering are shown, with the additive component marked by the shaded low-order modes and the cutoff indicated by the dashed line.

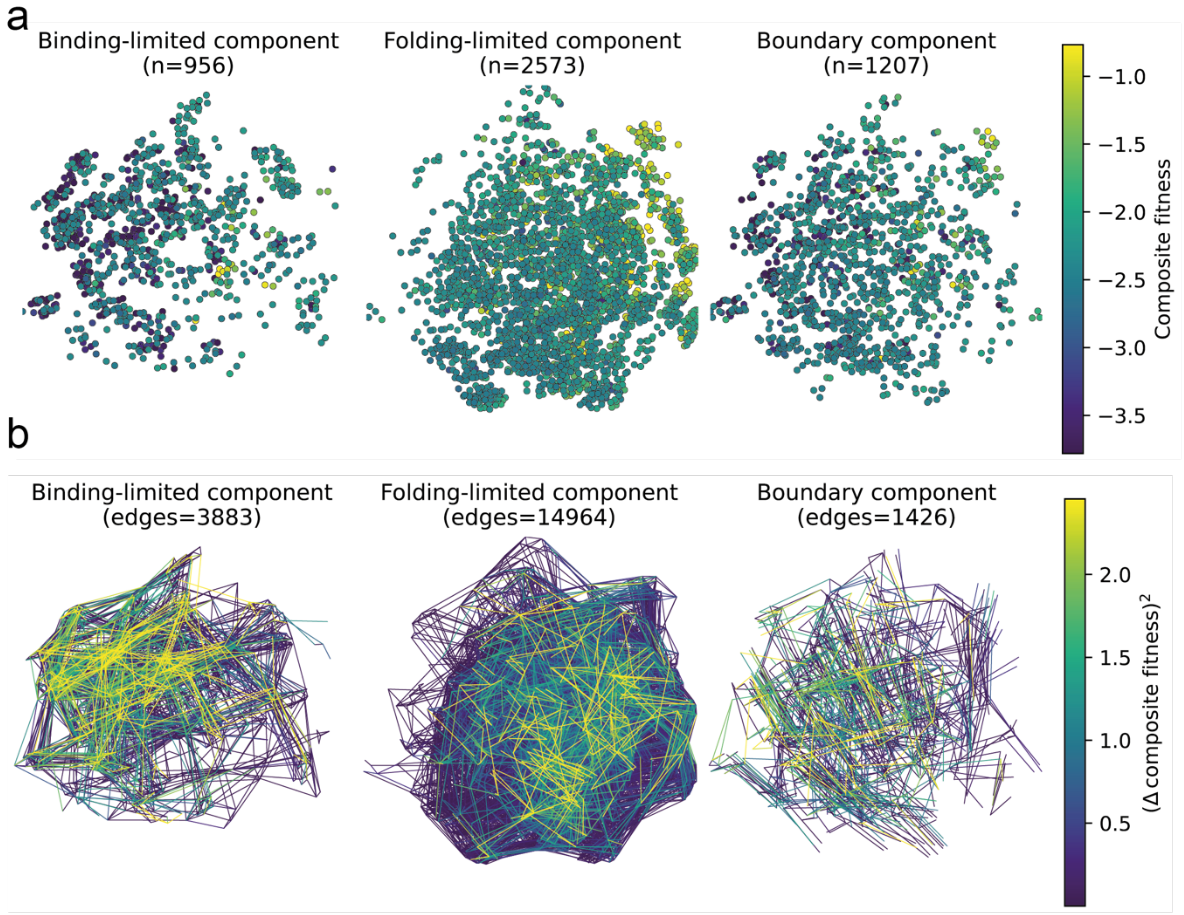


**SI Figure 18.** SH3 Hamming graph partitioned by phenotype regime. a. Binding-limited, folding-limited, and boundary components coloured by composite fitness. b. Edges in the same components coloured by squared composite-fitness change, showing where local Dirichlet energy is concentrated.

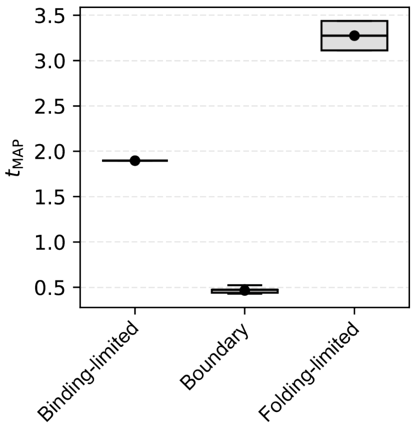


**SI Figure 19.** SH3 boundary ruggedness under uncertainty resampling. $t_{MAP}$ distributions for binding-limited, boundary, and folding-limited SH3 subgraphs after resampling latent folding and binding values within thermodynamic-model uncertainty.

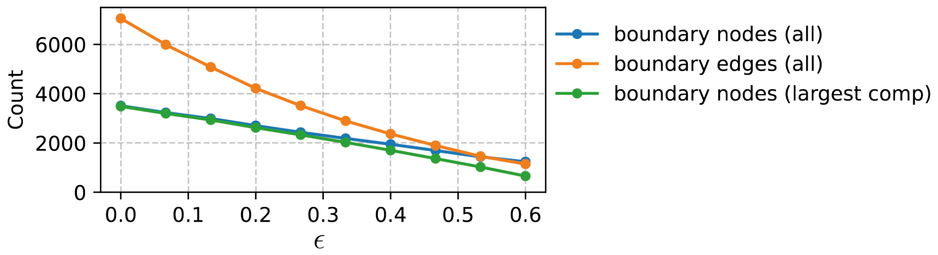


**SI Figure 20.** Boundary size as the limiter-index threshold ε increases. Counts of all boundary nodes, all boundary edges, and boundary nodes in the largest component decline as ε is tightened.

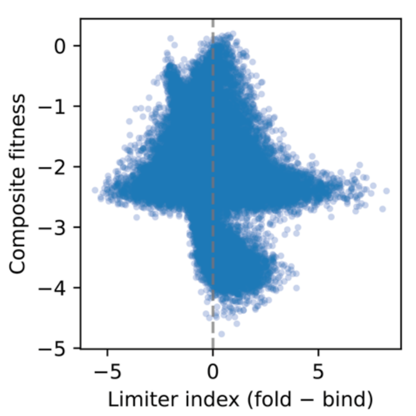


**SI Figure 21.** Relationship between limiter index and composite fitness in the SH3 landscape. Composite fitness is plotted against the folding-minus-binding limiter index; the boundary is centered near zero.

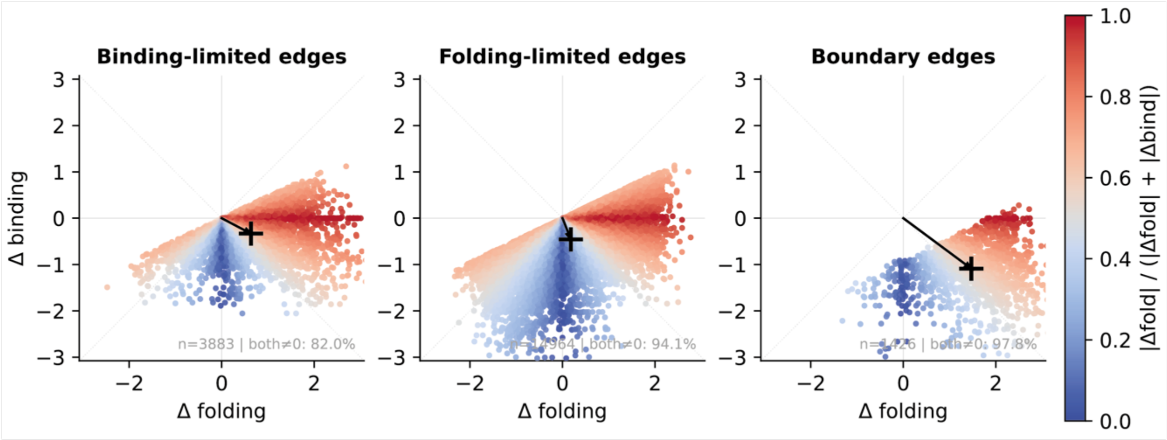


**SI Figure 22.** Vector differences in latent fitness contributions along SH3 edges. Edgewise changes in folding and binding contributions are shown for binding-limited, folding-limited, and boundary edges; colour indicates the fraction of total latent change attributable to folding.

**SI Text 1.** Lower bound on the Dirichlet energy, $E(f)$, by the number of edges that cross the cutset of $S$.

Consider the set of edges that connect nodes $i \in S$ and $j \in \overline{S}$, which we denote $\Delta S$,

$$\Delta S = \{(i,j) \in e \mid i \in S \ \wedge \ j \in \overline{S}\},$$

the minimum difference in fitness of any two nodes separated by $(i,j) \in \Delta S$, denoted $\delta$:

$$\delta = \min_{(i,j)\in\Delta S} [f_i - f_j] > 0,$$

and the definition of the Dirichlet energy as a sum of pairwise differences in signal:

$$E(\mathbf{f}) = \frac{1}{2} \sum_{(i,j)\in e} w_{i,j} [f_i - f_j]^2,$$

Where $w_{i,j}$ is the edge weight between incident nodes $i$ and $j$ with fitnesses $f_i$ and $f_j$. As $\Delta S \subset e$ and the Dirichlet energy is summed over all edges $(i,j) \in e$:

$$\frac{1}{2} \sum_{(i,j)\in\Delta S} w_{i,j} [f_i - f_j]^2 \le \frac{1}{2} \sum_{(i,j)\in e} w_{i,j} [f_i - f_j]^2.$$

Because $(i,j) \in \Delta S$ and by the definition of $\delta$ as the minimum difference in fitness between adjacent nodes connected by $(i,j) \in \Delta S$:

$$\delta^2 \le [f_i - f_j]^2.$$

Thus

$$\frac{1}{2} \sum_{(i,j)\in\Delta S} w_{i,j}\, \delta^2 \le \frac{1}{2} \sum_{(i,j)\in\Delta S} w_{i,j} [f_i - f_j]^2,$$

it follows that

$$\therefore \frac{1}{2} \delta^2 \sum_{(i,j)\in\Delta S} w_{i,j} \le E(\mathbf{f}).$$
